# Verkko: telomere-to-telomere assembly of diploid chromosomes

**DOI:** 10.1101/2022.06.24.497523

**Authors:** Mikko Rautiainen, Sergey Nurk, Brian P. Walenz, Glennis A. Logsdon, David Porubsky, Arang Rhie, Evan E. Eichler, Adam M. Phillippy, Sergey Koren

**Affiliations:** Genome Informatics Section, Computational and Statistical Genomics Branch, National Human Genome Research Institute, National Institutes of Health, Bethesda, MD USA; Department of Genome Sciences, University of Washington School of Medicine, Seattle, WA, USA; Howard Hughes Medical Institute, University of Washington, Seattle, WA, USA

## Abstract

The Telomere-to-Telomere consortium recently assembled the first truly complete sequence of a human genome. To resolve the most complex repeats, this project relied on manual integration of ultra-long Oxford Nanopore sequencing reads with a high-resolution assembly graph built from long, accurate PacBio HiFi reads. We have improved and automated this strategy in Verkko, an iterative, graph-based pipeline for assembling complete, diploid genomes. Verkko begins with a multiplex de Bruijn graph built from long, accurate reads and progressively simplifies this graph via the integration of ultra-long reads and haplotype-specific markers. The result is a phased, diploid assembly of both haplotypes, with many chromosomes automatically assembled from telomere to telomere. Running Verkko on the HG002 human genome resulted in 20 of 46 diploid chromosomes assembled without gaps at 99.9997% accuracy. The complete assembly of diploid genomes is a critical step towards the construction of comprehensive pangenome databases and chromosome-scale comparative genomics.

## Introduction

Recent advances in sequencing technologies have greatly increased the accuracy and length of sequencing reads ^1^. Pacific Biosciences’ high-fidelity (HiFi) reads can achieve accuracies of over 99.9% with read lengths of 18–25 kilobases (kb) ^2^ and Oxford Nanopore Technologies (ONT) reads routinely reach median lengths of 50–150 kb with accuracies around 95% ^3,4^. Recently, ONT has demonstrated the ability to generate relatively shorter reads (median 25–35 kb) at 99.9% accuracy. For convenience when not referring to a specific technology, we will refer to “long, accurate reads” (LA reads) as those with lengths greater than 10 kb and accuracy greater than 99.9%, and “ultra-long reads” (UL reads) as those with lengths over 100 kb and accuracies over 90%.

These technological advances have greatly simplified the process of reconstructing a genome from overlapping sequencing reads ^5,6^, yielding highly continuous genome assemblies ^4,7–11^. With the recent completion of the CHM13 human reference genome, the Telomere-to-Telomere (T2T) consortium demonstrated that gapless and accurate assembly of human genomes is now possible ^12-14^. However, this consortium effort required considerable resources, and the complete assembly of human chromosomes is not yet routine. While careful assembly of LA reads can complete a handful of chromosomes in a haploid human genome ^15^, the majority of chromosomes remain unresolved and require manual intervention ^11-13^. Chromosome-scale scaffolding of LA-based assemblies requires additional technologies such as Bionano ^16^, Strand-seq ^17-19^, or Hi-C ^20-22^, which can be error-prone ^11^ and require careful curation and validation ^23^.

Genome assembly is complicated due to the presence of large, highly similar repeats. The resolution of repeats during assembly requires either UL reads, to span repeat instances, or LA reads, to identify repeat-specific variants. UL reads can span exact repeats but lack the necessary accuracy to resolve long, very similar repeats. In contrast, LA reads excel at diverged repeats but fail on exact repeats, such as recent tandem duplications. Current methods for sequencing LA reads, such as PacBio HiFi, also suffer from known systematic sequencing errors ^8^, leading to regions of the genome with no coverage. On diploid genomes, LA reads can be used to build high-resolution haplotype-resolved graphs ^9,11^, where variants between haplotypes are represented as separate graph nodes (unitigs) with high accuracy and not collapsed into a mosaic assembly ^8,9,11^. However, their relatively short read lengths limit phasing to a few hundred kb in human genomes. As a result, most LA assemblers output pseudo-haplotypes, a random walk combining the short, but haplotype-resolved, unitigs into long, but haplotype-mixed, contigs. In contrast, UL reads can phase over megabases ^24^ but have lacked the accuracy to separate haplotypes within the assembly graph or generate a high-quality consensus sequence ^4^. Thus, LA and UL sequencing reads are complementary for genome assembly but no tools exist for co-assembling them.

Here, we present our new assembler, Verkko, which combines the best features of LA and UL reads into one workflow. Previous hybrid assembly approaches have relied on short reads to build the initial graph and were limited to prokaryotic genomes ^25-27^. Verkko was specifically designed for large, eukaryotic genomes and incorporates LA reads, UL reads, and haplotype information from familial trios ^28^, Hi-C ^20,21^, or Strand-seq ^29,30^ for the complete assembly of diploid chromosomes.

## Results

### Verkko overview

Verkko builds on lessons learned from the completion of the CHM13 human reference genome and adopts a similar graph-first approach. However, the methods developed for CHM13 were semi-manual and not directly applicable to heterozygous genomes ^14^. Verkko extends this strategy to diploid genomes and fully automates the process. Conceptually, Verkko constructs a high-resolution assembly graph from LA reads that resolves diverged repeats and captures small variants between the repeats and haplotypes. This graph is then progressively simplified by the integration of UL reads and haplotype markers to bridge exact repeats, fill coverage gaps, and phase haplotypes (Fig. 1) (Online Methods).

**Figure 1.**
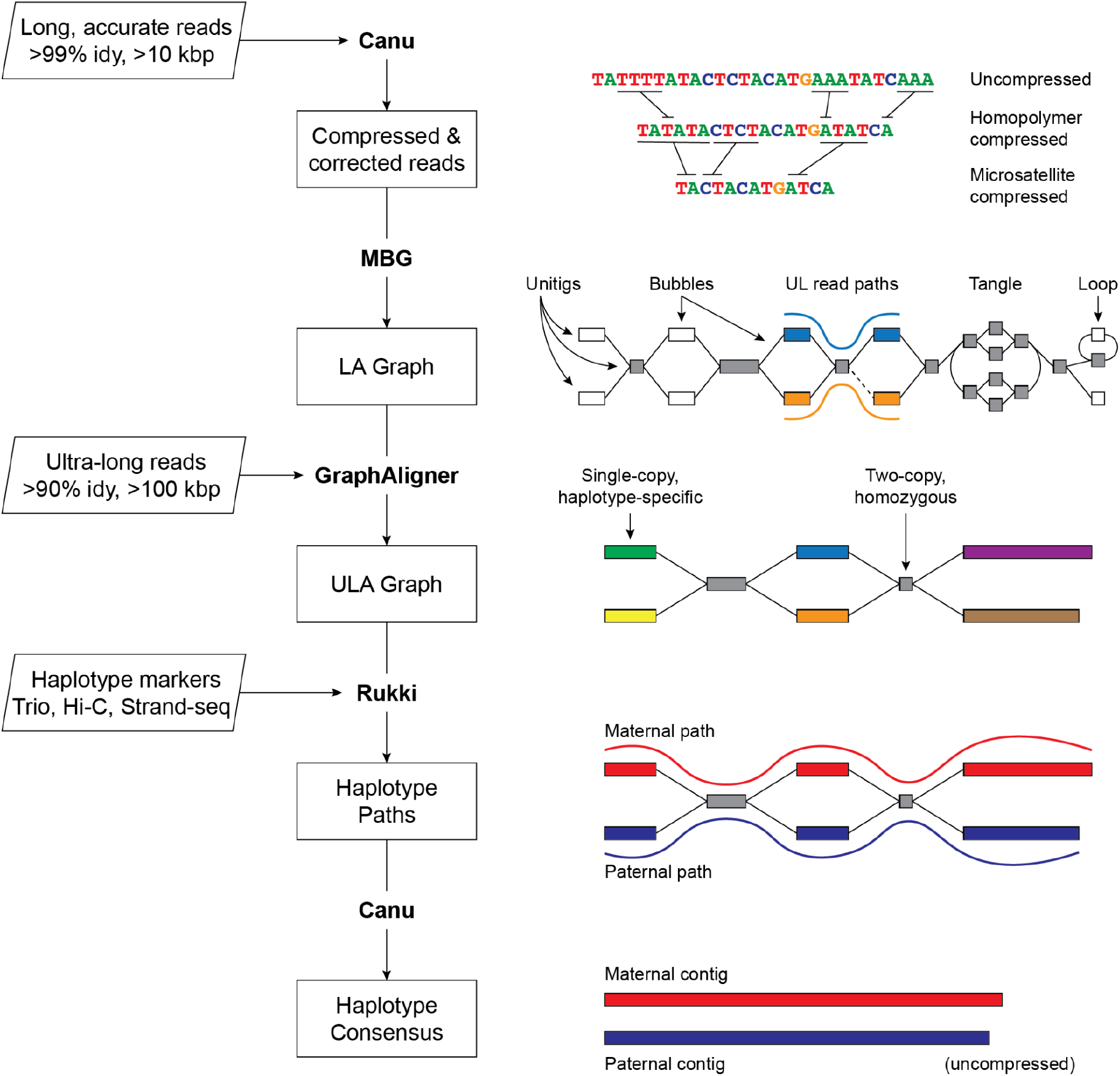
Verkko assembly workflow. Inputs are on the left, with program outputs listed in rectangles. Key components of the pipeline are highlighted, including Canu ^8^, MBG ^33^, GraphAligner ^34^, and Rukki (Online Methods). Using the outputs of these tools, Verkko performs successive rounds of processing and graph resolution. Input reads are first homopolymer compressed and error corrected, and remain so throughout the process. MBG also compresses microsatellites to aid graph construction, but only internally. The first graph output by Verkko is an accurate, high-resolution de Bruijn graph built from the long, accurate reads (LA Graph). This is a node-labeled graph comprising unitigs (nodes) and their adjacency relationships (edges). Ultra-long reads are then aligned to the LA graph to identify read paths (blue curve, orange curve), and Verkko uses them for phasing bubbles, filling gaps (dotted edge), and resolving loops and tangles. The resulting simplified graph (ULA Graph) is typically composed of single-copy, haplotype specific unitigs, separated by two-copy, homozygous unitigs that could not be phased from the read data alone. Haplotype-specific markers are then used to label nodes in the ULA graph and identify haplotype paths through the graph (maternal, red; paternal, blue). These paths are converted to haplotype-specific contigs using a consensus algorithm that recovers the homopolymers.

Homopolymer insertions and deletions are one of the primary sources of error in long-read sequencing, and compressing them (e.g. A_1_…A_*n*_ becomes A_1_ for all *n* > 1) simplifies the assembly process ^8,14,31^. The entire Verkko pipeline operates on homopolymer-compressed sequences, which are recovered during the final consensus phase. After compression, the LA reads are error-corrected and used to build a multiplex de Bruijn graph ^15,32^. UL reads are then aligned to this graph to patch coverage gaps and further resolve repeats and haplotypes. The graph is finally cleaned, and haplotype paths are identified using haplotype-specific markers from additional parental or long-range sequencing information. To restore homopolymers, LA read paths from the initial graph are lifted to the final graph and a consensus sequence is computed for all nodes and haplotype paths.

Verkko’s final output is a phased, diploid assembly of both paternal and maternal haplotypes, as well as a highly accurate and resolved assembly graph that can assist with additional genome finishing. In cases where the available data could not fully resolve a chromosome as a single contig, Verkko leverages the assembly graph structure to generate telomere-to-telomere, haplotype-resolved scaffolds, thus removing the need for a separate, error-prone scaffolding step ^11,23^.

### Complete haploid genome assembly

We tested Verkko (version beta2) (Code availability) on a recently published *Arabidopsis thaliana* genome ^35^ (Data availability), and benchmarked it against state-of-the art assemblers designed for HiFi ^9,15,36^ and ONT ^7^ data (Supplementary Note 1, Supplementary Table 1). We compared the assemblers to the published *A. thaliana* reference ^35^ using Quast ^37^ (Supplementary Note 2). The errors were filtered to exclude those arising from likely mis-assemblies in the reference. Using only the HiFi data, Verkko was comparable to other HiFi-only assemblers, failing to resolve any chromosomes end-to-end. However, when combining HiFi and ONT data, Verkko assembled 4 of 5 chromosomes into single unitigs spanning >99.5% of the reference genome, with 2 of the 4 having canonical vertebrate telomeric repeats on both ends (Supplementary Table 2). The only exception was Chr4, which had a single heterozygous bubble and unresolved 45S rDNA tangle. Verkko also had the lowest error count and comparable base accuracy to other HiFi-based assemblers (Supplementary Table 2). All assemblies had a lower number of differences when compared to the Verkko assembly than the published reference, suggesting that the Verkko assembly was more correct (Supplementary Table 3). This finding was further supported by a reference-free evaluation using VerityMap ^38^ (Supplementary Figs. 1–5).

The T2T consortium relied on an almost fully homozygous cell line for completion of the human genome ^14^. Due to its completeness, extensive validation, and cataloged variants ^39^, the CHM13 cell line provides an excellent test case for benchmarking the assembly of a single haplotype. We ran Verkko with the same HiFi and ONT data that was originally used to assemble the CHM13 genome (Data availability), and compared to HiFi-only and ONT-only assemblers (Table 1, Fig. 2). The assemblies were validated with QUAST ^37^ and VerityMap ^38^ while filtering for known heterozygous variants. Verkko correctly assembled 12 chromosomes from telomere to telomere, with 5 additional chromosomes assembled into a single unitig containing >95% of the expected sequence. This is double that of any assembly based on a single technology, with the next best result coming from LJA ^15^, which completely assembled 6 chromosomes from HiFi data alone.

**Table 1.**
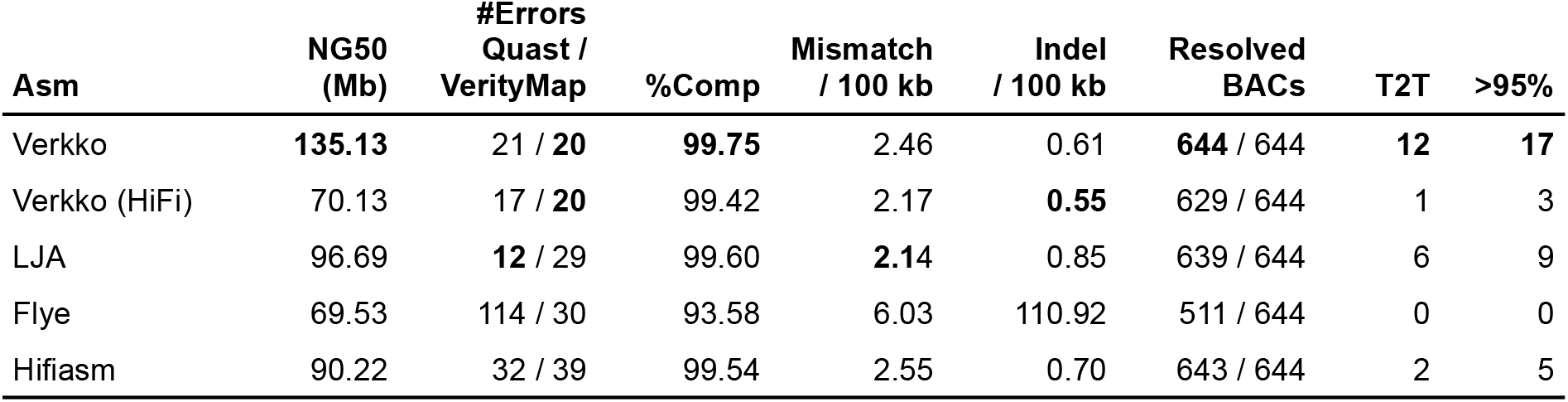
Validation of CHM13 assemblies. We used the published reference of CHM13v1.1 ^14^ to evaluate the assemblies with QUAST and ran VerityMap as a reference-free alternative. The relative assembler ranking between QUAST and VerityMap shows good agreement, only switching LJA and Verkko in terms of fewest errors. %Comp: chromosome completeness as measured by QUAST alignments. Number of mismatches and Indels reported by QUAST are given per 100 kb. T2T: reports complete chromosomes with a single unitig covering >99% of the reference bases and having canonical telomeres on each end. >95%: reports chromosomes with a single unitig covering >95% of the reference with no requirement for telomeres. QUAST errors intersecting known heterozygous variants or errors ^39^, as well as those within the core rDNA arrays, were excluded using a filter script from Shafin *et al*. ^4^. CHM13 reference BACs were evaluated as in Nurk *et al*. ^8^. Canonical telomeres were identified using the VGP pipeline ^10^.

| Asm | NG50 (Mb) | #Errors<br>Quast /<br>VerityMap | %Comp | Mismatch<br>/ 100 kb | Indel<br>/ 100 kb | Resolved<br>BACs | T2T | >95% |
| --- | --- | --- | --- | --- | --- | --- | --- | --- |
| Verkko | **135.13** | 21 / **20** | **99.75** | 2.46 | 0.61 | **644** / 644 | **12** | **17** |
| Verkko (HiFi) | 70.13 | 17 / **20** | 99.42 | 2.17 | **0.55** | 629 / 644 | 1 | 3 |
| LJA | 96.69 | **12** / 29 | 99.60 | **2.14** | 0.85 | 639 / 644 | 6 | 9 |
| Flye | 69.53 | 114 / 30 | 93.58 | 6.03 | 110.92 | 511 / 644 | 0 | 0 |
| Hifiasm | 90.22 | 32 / 39 | 99.54 | 2.55 | 0.70 | 643 / 644 | 2 | 5 |

**Figure 2.**
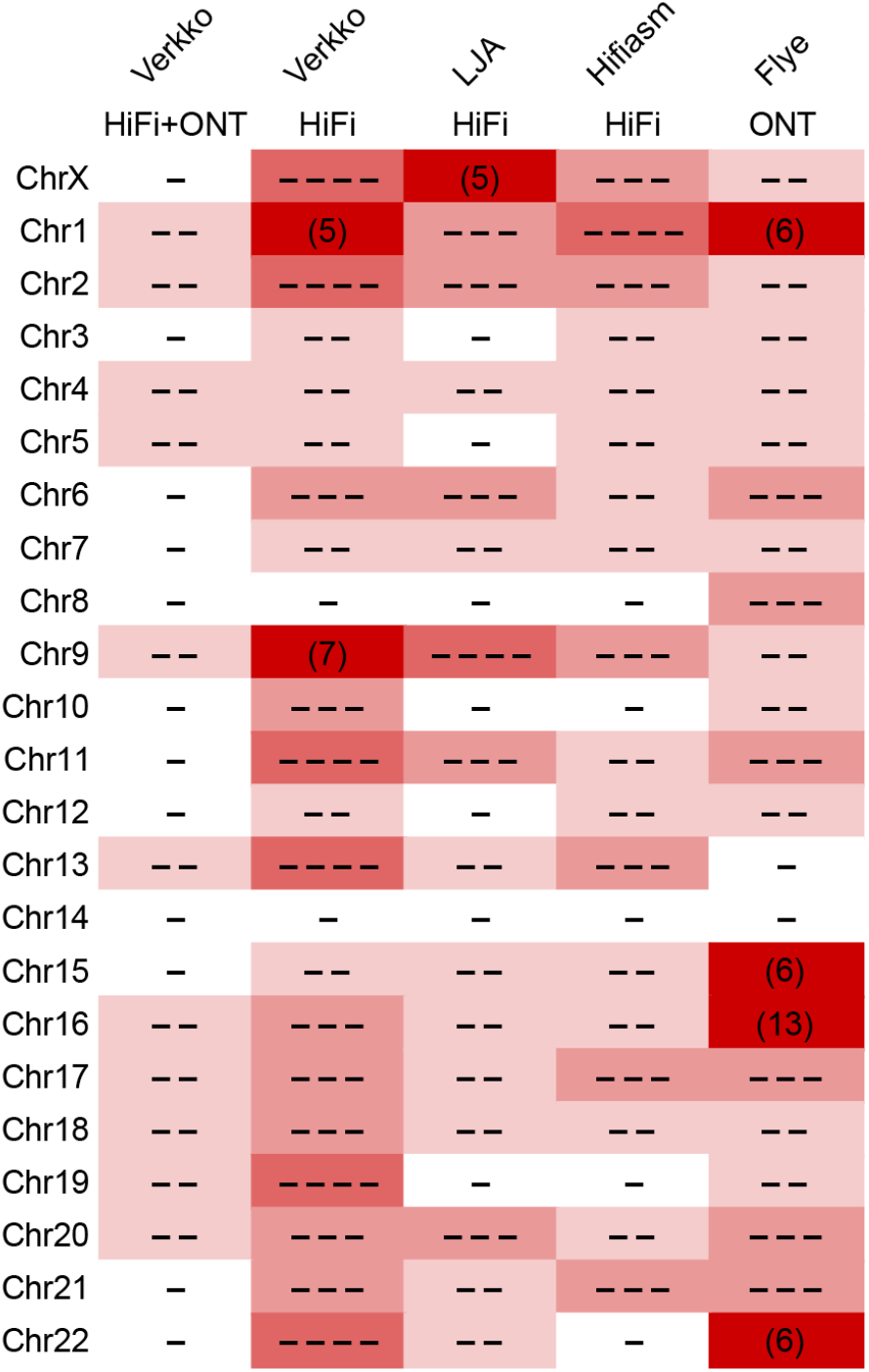
Continuity of assembled CHM13 chromosomes. The minimum number of continuous alignments needed to cover 85% of each chromosome (LGA85) is represented by the number of dashes, or plain numbers when :25. Higher numbers are shaded progressively darker. Alignments were taken from QUAST output versus the CHM13v1.1 reference, after filtering errors at heterozygous sites and potential erroneous regions in the reference as in Table 1. Verkko using HiFi data alone is comparable to other HiFi-based assemblies ^9,15^, but Verkko combining HiFi and ONT data has the most complete and correct chromosomes (12).

Even in cases where Verkko did not assemble a complete chromosome, the assemblies were close to complete and the correct resolution was often clear from the final graph (Supplementary Fig. 6). Unresolved regions were limited to the highly repetitive rDNA arrays and centromeric human satellite arrays on Chr9 and Chr16. However, even the rDNA regions were partially resolved, correctly separating Chr15 and Chr22 into their own connected components. The component containing Chr13, Chr14, and Chr21 had three separate rDNA tangles, with the distal and proximal regions of each chromosome connected to the corresponding rDNA tangle (Supplementary Fig. 7). It was previously noted that the these chromosomes shared the highest degree of inter-chromosomal similarity in the human genome ^14^, which may explain why they are the only CHM13 chromosomes not separated into distinct components of the graph.

Verkko’s results show the advantage of combining LA and UL data types. For example, Chr8 was >90% complete in LJA ^15^, Hifiasm ^9^, and HiFi-only Verkko. However, there is a coverage gap at the end of this chromosome due to a GA-rich microsatellite that requires ONT-based gap-filling to resolve ^8,13^. Verkko correctly identified the missing component and patched this gap. Other chromosomes, such as ChrX, are specifically enriched for HiFi coverage gaps and large near-perfect repeats ^8^. No HiFi-only assembly recovered more than 40% of ChrX in a single contig, and at least 7 contigs were needed to cover >99% of it. Flye’s ^7^ ONT-based assembly was more continuous but failed to resolve the centromeric repeats. In contrast, a single unitig in Verkko contained >99.98% of ChrX, missing only the last 50 kb on the q-arm due to a retained heterozygous bubble. We manually inspected the VerityMap output for the Verkko assembly and found that some reported QUAST errors may be false positives or errors in the CHM13 reference, as there is better agreement with the Verkko assembly in these regions (Supplementary Figs. 8 and 9 for Chr4 and Chr17, respectively).

### Complete diploid genome assembly

To evaluate Verkko on a highly heterozygous sample, we chose the Darwin Tree of Life insect genome *Harmonia axyridis* ^40^ (Data availability). Since Verkko outputs phased unitigs, we compared its output to fully phased equivalents from other assemblers. Despite relatively short ONT data (<2x sequence read coverage ≥2100 kb), Verkko’s unitig NG50 was 14.53 Mb, similar to the pseudo-haplotype reference NG50 of 15.11 Mb and much larger than the phased unitigs produced by HiFi-only assemblers: Verkko (HiFi) (6.17 Mb), LJA (5.24 Mb), or Hifiasm (7.08 Mb). We did not evaluate the quality of the assemblies with QUAST as the reference likely contains errors, but Verkko had a low number of errors (18) identified by VerityMap ^38^. Thus, Verkko’s combination of HiFi and ONT sequencing not only produces highly accurate and complete assemblies but can produce phased unitigs rivaling the best unphased pseudo-haplotypes for diverse genomes. When incorporating Hi-C data ^41^, Verkko’s contig NG50 increased to 25.31 Mb, far exceeding the continuity of the pseudo-haplotype reference.

We ran Verkko on the benchmark HG002 human sample from Genome in a Bottle (GIAB) ^42^ and the Human Pangenome Reference Consortium (HPRC) ^43^ (Data availability). We used the downsampled 35× dataset base-called with DeepConsensus, which has been shown to improve coverage and assembly quality ^44^, along with 60× ONT ultra-long reads. We compared the results to the recently finished HG002 ChrX and ChrY ^14^ using QUAST and evaluated assembly quality and accuracy using reference-free methods (Supplementary Note 2). We layered Hi-C and trio information (Supplementary Note 1) onto the Verkko graph and compared it to other state-of-the-art phased assembly results ^9,36^ (Table 2, Supplementary Table 4).

**Table 2.** Quality and completeness of HG002 diploid assemblies. We evaluated contig NG50 and phase accuracy for all assemblies using Merqury ^45^. With a goal of phased assemblies, pseudo-haplotype outputs from Flye and Hifiasm were not considered. We evaluated hamming error rate (the fraction of non-dominant allele variants in a unitig) and Phred QV ^46^ using Merqury. We measured errors versus the recently finished HG002 chromosomes X and Y ^14^ using QUAST. Results are shown for DeepConsensus HiFi ; results from the original HiFi reads are in Supplementary Table 4. LJA and Hifiasm do not output scaffolds, so no scaffold NG50 values are reported. Verkko assemblies used 105x HiFi coverage (mean 17.5 kbp, Data availability) while the HPRC curated assembly used 130x HiFi coverage (mean 14.8 kbp) ^11^.

| Asm | Contig<br>NG50 (Mb) | Scaffold<br>NG50 (Mb) | Hamming<br>error | QV | #Errors<br>Chr X & Y |
| --- | --- | --- | --- | --- | --- |
| **Downsampled (35× HiFi / 60× ONT-UL)** |  |  |  |  |  |
| Verkko | 12.90 |  | 0.13% | 52.18 | 10 |
| Verkko + trio | **80.77** | **102.55** | 0.13% | 52.40 | 10 |
| Verkko + Hi-C | 58.24 | 82.42 | 0.16% | 52.48 | 10 |
| LJA | 0.39 |  | 0.14% | 55.75 | 7 |
| Hifiasm (unitigs) | 0.35 |  | 0.23% | **61.05** | **2** |
| Hifiasm + trio | 64.50 |  | **0.06%** | 60.86 | 26 |
| Hifiasm + Hi-C | 66.34 |  | 0.79% | 60.57 | 37 |
| **Full-coverage (>100× HiFi / 85× ONT-UL)** |  |  |  |  |  |
| Verkko | 17.35 |  | **0.05%** | 54.91 | **5** |
| Verkko + trio | **134.00** | 135.80 | **0.05%** | 55.77 | **5** |
| Verkko + Hi-C | 68.32 | 85.97 | **0.05%** | 55.57 | **5** |
| HPRC curated | 72.70 | **146.75** | 0.13% | **61.35** | 26 |

Verkko produced megabase-scale phase blocks, similar to our observation on non-human genomes. We found that DeepConsensus HiFi increased NG50 without introducing errors and decreased the switch rate in most cases (Hifiasm + Hi-C has a higher switch rate). Verkko has a low switch rate and a low error count with a QV over 50 (1 error per 100 kb, Supplementary Figs. 10–12). When adding Hi-C or trio information, the Verkko assembly had fewer errors when compared to Hifiasm (3.7-fold less with Hi-C and 2.6-fold less with trios). Verkko has a higher switch error rate than Hifiasm using trios but a lower switch rate when using Hi-C. Our current integration of Hi-C or trio data does not correct switch errors in the initial assembly and, thus, cannot be below the 0.13% switch rate of the Verkko unitigs.

Both Verkko and Hifiasm assemblies are highly complete, with Verkko recovering slightly more multi-copy genes but at the expense of a slightly higher false-duplication rate within individual haplotypes (Supplementary Table 5). This effect was most evident when using Hi-C data due to the higher rate of haplotype misassignment or lack of assignment compared to trios. Post-processing tools, such as purge_dups ^47^, or improved Hi-C handling by Verkko can potentially address these duplicated sequences when trio information is unavailable.

Uniquely, Verkko is able to output accurate scaffolds using only the connectivity information contained within the assembly graph. For the downsampled HG002 dataset, this feature produced chromosome-scale scaffolds for 7 and 4 chromosomes with trio and Hi-C information, respectively. Previously, such continuity would have required a separate and error-prone scaffolding step ^11^.

Finally, we used the full-coverage HG002 dataset (105× HiFi, 85× ONT-UL) to produce the most continuous assembly of this genome to date (Fig. 3). We compared the Verkko assembly to a recently published benchmark assembly which used high-coverage HiFi data for contig assembly, ONT assembly for gap filling, trio information for phasing, BioNano as well as Hi-C data for scaffolding, and manual curation. Both the Verkko trio and Hi-C assemblies have fewer errors and more accurate phasing than the benchmark ^11^, and the trio-based Verkko assembly has nearly double the contig NG50 size (Table 2). The improvement in accuracy is partially due to the avoidance of large-scale switches within complex regions of the centromeric repeat arrays (Supplementary Fig. 13). In contrast to Hifiasm, Verkko was able to resolve these regions using the complementary ONT data, enabling more accurate phasing.

**Figure 3.**
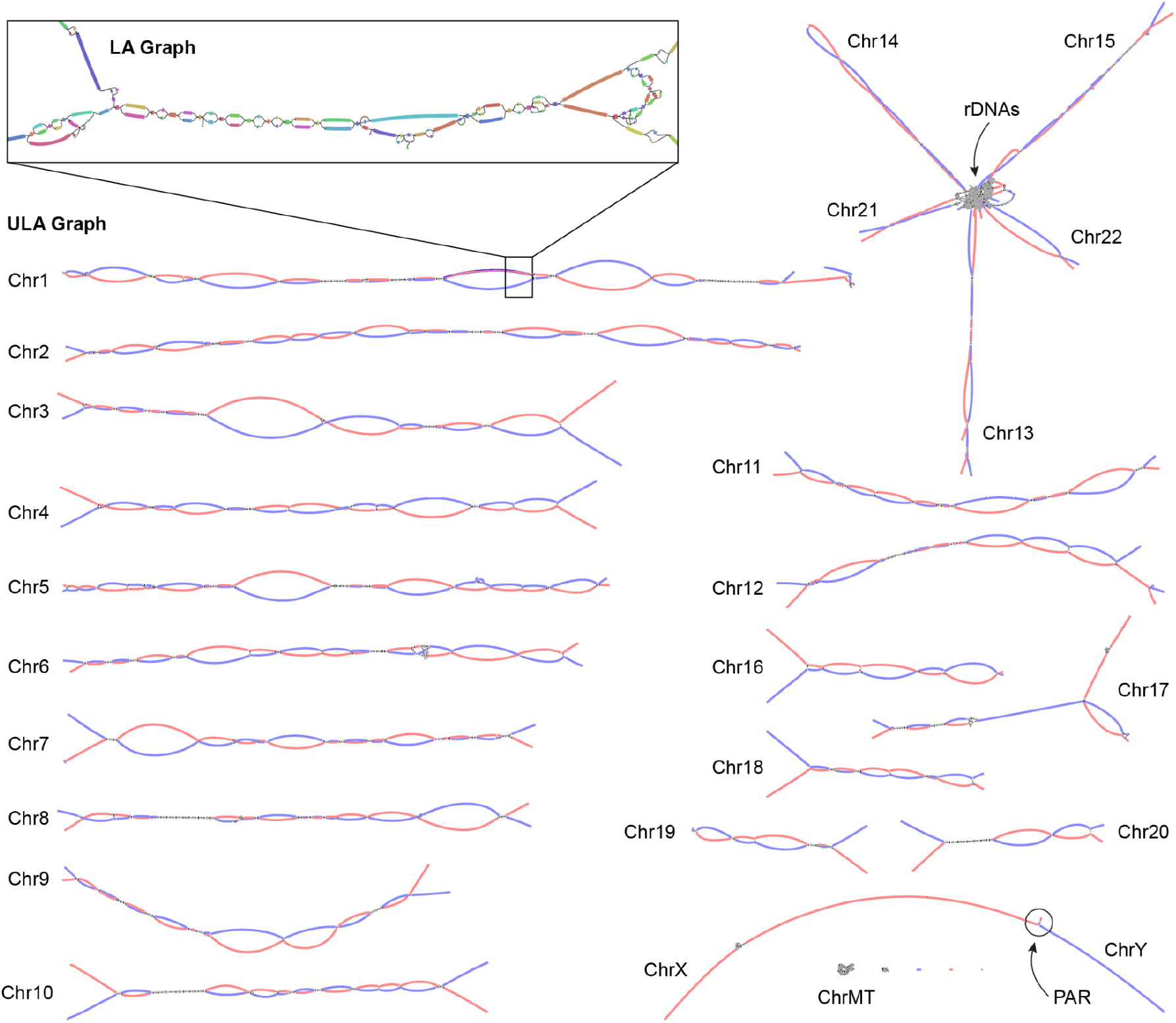
Verkko assembly graph of the HG002 diploid genome. ULA graph integrating both PacBio HiFi and ONT ultra-long reads. Inset shows a portion of the LA graph with multiple bubbles, tangles, and tips prior to resolution using the ultra-long reads. This particular region contains stretches of homozygosity and exact repeats that could not be resolved by HiFi alone. After integration of the ONT data, the corresponding region in the ULA graph is resolved into two linear haplotypes. Each chromosome in HG002 is mostly resolved as a single connected component, with the exception of the acrocentric (13, 14, 15, 21, and 22) and sex chromosomes (X and Y), which are joined by the highly similar rDNA arrays and pseudoautosomal regions (PAR), respectively. The final unitigs in the ULA graph have been colored after assembly by trio-derived haplotype markers (maternal, red; paternal, blue). Using this information to determine haplotype paths through the graph, Verkko was able to completely assemble 20 chromosomal haplotypes in HG002 from telomere to telomere without gaps.

Of the 46 chromosomes in HG002, Verkko resolves 27 of them into single scaffolds using the trio information. Of these, 20 are completely assembled from telomere to telomere without gaps, which is a dramatic improvement over the previous HG002 benchmark assembly ^11^ that includes 19 complete scaffolds and only one complete contig. Using the Hi-C data alone, Verkko was able to resolve 9 chromosomes into single scaffolds, 7 of which are gapless.

We evaluated the full-coverage Verkko trio assembly using orthogonal Strand-seq ^29,48^ data from the same sample, which allows for the detection of inversions and translocations in the assembly. We found that the Verkko trio assembly had a higher fraction of contigs and bases correctly assigned to chromosomes than the benchmark assembly ^11^ (Supplementary Fig. 14 and 15). The majority of chromosomes were covered by a single Verkko scaffold for both the maternal and paternal haplotypes, and there were no mis-orientations or regions which failed to resolve a homozygous inversion, again an improvement on the previous result. There were a handful of regions, accounting for approximately 30 Mb per haplotype, where there was an ambiguity, either due to true heterozygous inversions between the haplotypes or due to low mappability of Strand-seq data (Supplementary Fig. 16). After filtering ambiguous regions, we confirmed 18 heterozygous inversions ranging from 6 kb to 4.1 Mb (median 237 kb) and accounting for 9 Mb per haplotype (Supplementary Table 6, Supplementary Fig. 17). This includes three medically relevant regions which were not correctly resolved in previous HG002 assemblies ^11^. The largest of these is a known, medically relevant, recurrent inversion on Chr8 ^13,49^. As observed with CHM13, the Verkko reconstruction improves on the manually curated reference and was used to confirm an inversion error in the T2T assembly of ChrY (A. Rhie, personal communication) (Supplementary Fig. 18).

Verkko was able to completely resolve both haplotypes of a single chromosome in several cases (Fig. 4, Supplementary Figs. 19 and 20). This enabled us to assess variability between centromeric regions in a diploid human genome. Consistent with prior studies ^50-57^, we found that centromeric α-satellite higher-order repeat (HOR) arrays often vary in length by tens to hundreds of kilobases. For example, Figure 4 shows the centromeric HOR arrays from Chr19. Both Chr19 haplotypes are well-supported by read alignments, indicating no large-scale mis-assemblies. The arrays differ in length with respect to different individual HOR units, suggesting distinct patterns of HOR expansion between the maternal and paternal haplotypes. The paternal haplotype’s D19Z3 array is approximately 620 kb larger than the maternal. The D19Z3 array also differs in HOR structure between the haplotypes. While both arrays are primarily composed of a 2-mer, the paternal haplotype is flanked by 4-mer and 8-mer HORs and the maternal haplotype is dominated by the 2-mer HOR. The HOR arrays show evidence of layered expansions ^13^, with more recently evolved HOR repeats that have expanded within the core of the arrays (shown in dark orange) and, consequently, pushing more divergent HOR arrays to the sides (shown in green to yellow). The other 8 pairs of completely assembled centromeric satellite arrays (chromosomes 1, 3, 4, 9, 10, 11, 16, and 18) show similar patterns of differential expansion, contraction, and homogenization, underscoring the ubiquity of centromere HOR array size variation between haplotypes (Supplementary Figs. 21 and 22).

**Figure 4.**
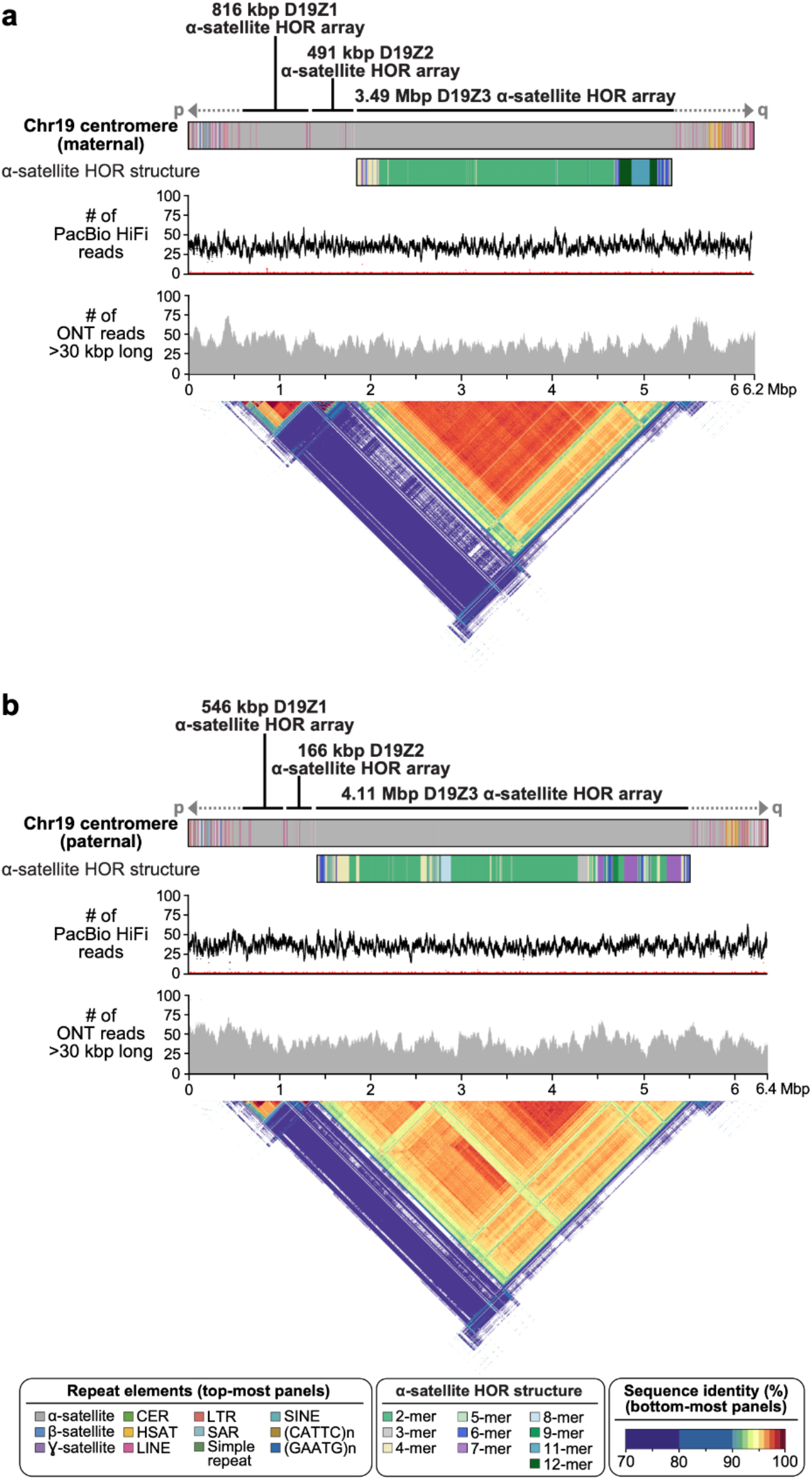
Comparison of the maternal (a) and paternal (b) Chr19 centromeric regions in HG002. Each haplotype is annotated with its repeat structure (top-most track), HOR structure, PacBio HiFi coverage, and ONT coverage. Last, the triangle shows sequence similarity within each haplotype ^58^. Both haplotypes reveal the presence of three higher-order repeat (HOR) arrays (D19Z1, D19Z2, and D19Z3) that vary in size and structure.

## Discussion

Verkko is a new assembler that leverages the complementarity of long, accurate reads and ultra-long reads to produce assemblies that are more continuous and accurate than when using either technology alone. Verkko relies on high-accuracy reads to build an initial assembly graph which is further resolved with long, noisy reads. Haplotype markers can then be mapped to identify haplotype paths through the graph. For complete, diploid genome assembly, we currently recommend sequencing approximately 50× genomic coverage of LA reads, 50× in UL reads >100 kb, and 50× of parental short reads. High-quality draft assemblies are possible with lower coverage, and Hi-C or Strand-seq can be used for chromosome-scale phasing in the absence of trios.

Vekko is a modular pipeline that is adaptable to different technologies or the substitution of specific components. For example, Verkko should be compatible with other LA read technologies, such as ONT “duplex” sequencing ^59^. Additionally, the graphs produced by Verkko are similar in spirit to those from LJA ^15^, so LJA graphs could potentially be used as a basis instead of MBG graphs. However, Verkko’s methods for UL resolution and haplotype walking were developed using HiFi-based MBG graphs and would likely require tuning for the idiosyncrasies of different data types and tools.

On haploid genomes, Verkko automates telomere-to-telomere assembly for the majority of human chromosomes. On diploid genomes, Verkko generates megabase-scale phase blocks, rivaling the continuity of pseudo-haplotypes single-technology assemblers on non-human genomes. In combination with Hi-C or trio information, Verkko can generate complete, haplotype-resolved scaffolds for a subset of chromosomes given sufficient sequencing coverage.

Verkko’s current Hi-C integration requires pre-binned haplotype reads or assemblies, such as those generated by DipAsm ^41^ or Hifiasm ^36^. However, it would be possible and likely more accurate to infer haplotype walks from Hi-C reads aligned directly to the graph ^36^. Another limitation of our current approach is that trio or Hi-C information is used after LA and UL read resolution. As a result, switch errors introduced during graph construction are not corrected by later stages. To address this, future versions could incorporate haplotype information in tandem with graph construction, or could break erroneous graph nodes based on haplotype markers. Despite this limitation, we found haplotype switches to be rare in practice, with <1% of the assembly in nodes with :25% hamming error in the downsampled HG002 dataset, and <0.3% in the full-coverage dataset.

With accurate and phased assemblies of complete human chromosomes, we observed large-scale variation between the most repetitive regions of human haplotypes, with the centromeres having an elevated level of heterozygosity compared to the rest of the genome (Supplementary Figs. 19 and 20). The autosomal centromeric arrays show differentiated expansion of HOR elements and differ in their HOR copy age. This is in contrast to the high similarity previously observed between ChrX centromeric arrays ^60^ but agrees with a previous analysis of Chr8 ^13^. Our observation is consistent with a low rate of recombination within the centromeric arrays and supports the idea of centromere haplogroups ^61^.

With its ability to resolve complete haplotypes, Verkko ushers in a new era of comprehensive genomic analysis. The complete assembly of vertebrate haplotypes has direct application for the construction of new reference genomes, and ultimately, to better understanding of the relationships between large, complex structural variation, phenotype, and disease.

## Online Methods

Verkko (“network” or “graph” in Finnish) includes a core set of tools, including HiCanu ^8^, MBG ^33^, and GraphAligner ^34^, that have been extended and integrated together for the assembly of complete haplotypes. All steps of the pipeline are described below along with the details of any improvements made to the existing tools since their published versions.

Although “sequencing coverage” traditionally means the total bases sequenced divided by the haploid genome size, this section considers the diploid genome. Thus, 50× genomic coverage equates to roughly 25× coverage per haplotype. Similarly, when discussing “single-copy” or “unique” nodes within the assembly graph, this is again in regard to the diploid genome. A typical node-labeled diploid assembly graph comprises single-copy nodes that are haplotype-specific and multi-copy nodes that represent homozygous sequences between the two haplotypes (typically two-copy) or genomic repeats (two or more copies).

Verkko’s goal is to resolve each chromosome into one single-copy node spanning from telomere to telomere. However, early iterations of the assembly graph usually contain a number of “bubbles” and “tangles”. A bubble is a subgraph between source and sink nodes *v* and *w*, where only *v* and *w* have connections to nodes outside the bubble. Heterozygous variants are a common cause of simple bubbles, where nodes *v* and *w* are both two-copy, homozygous nodes connected by exactly two single-copy, disjoint paths representing the distinct haplotypes. Somatic variants and uncorrected sequencing errors are another source of bubbles, and can be identified by their low coverage. Lastly, a tangle is a subgraph containing multi-copy nodes and whose nodes only connect to other non-unique nodes within the tangle or to unique nodes outside the tangle. Tangles are caused by long, exact or near-exact repeats either within or between haplotypes.

### Error correction and homopolymer compression

Correction of the LA reads follows the procedure described in HiCanu ^8^ with incremental improvements to the accuracy. First, to mask a primary source of errors in LA reads such as PacBio HiFi, all homopolymers are compressed to a single base. The reads are then aligned in an all-vs-all manner and compared. If a read has a position which is covered by multiple aligned reads and most other reads agree on a difference at that position, the position in the read is considered to be erroneous and corrected. If at least two other reads support the base, it is left unchanged. The corrected, homopolymer-compressed LA reads are used for all downstream stages of the pipeline, and only reverted to their original form during the final consensus stage.

### Microsatellite compression

Although homopolymer errors are the most common systematic error type in HiFi reads ^2,8^, they are not the only error type. Microsatellite errors happen when a read contains a short, exact, tandem repeat with some copies incorrectly deleted or inserted. For example, the genomic sequence ACGACGACG, composed of the unit ACG repeating three times, could be miscalled as ACGACGACGACG, containing one extra unit copy. We have extended MBG to perform microsatellite compression to mask these errors.

The microsatellite compression used by MBG works analogously to, and follows directly after, homopolymer compression. Each microsatellite repeat unit is represented by a unique character, and any tandemly repeating characters are then merged together as with homopolymers. MBG transforms the input read into a special alphabet where each possible microsatellite repeat with a unit size of up to 6 bp gets its own character. Microsatellites are detected in the reads whenever a unit repeats at least twice and are encoded by three properties: the unit sequence, unit length, and overhang length. The overhang length is always strictly less than the unit length and represents a partial unit at the end of the repeat. The overhang sequence does not need to be encoded since it is always a prefix of the unit sequence. This encoding considers repeats with different overhangs to be distinct characters and does not include any information regarding copy number. Given a unit length *n*, there are *n*4^*n*^ possible microsatellite characters. Considering all possible units of size 1 to 6 nucleotides, there are less than 2^16^ characters in the alphabet and so each can be represented by a 16-bit integer.

Reverse complements are handled by computing the reverse complement of the unit sequence and rotating it according to the overhang length. Here we represent the encoding with the notation (*x*)*y*, where *x* is the unit sequence and *y* is the overhang sequence. For example, the repeat ACGACGACGAC is written as (ACG)AC, with a unit length 3, unit sequence ACG, and an overhang length of 2. The reverse complement of a character can be found in this notation by reverse complementing the string and keeping the parenthesis in place. In this example, the reverse complement of (ACG)AC is (GTC)GT.

Some microsatellite repeats can be encoded in multiple ways. For example, the sequence ATATATATATATAT could be encoded as (AT), (ATAT)AT, or (ATATAT)AT. If there are multiple possible encodings, we pick the one with the shortest unit length. Also, any microsatellite repeat contained within another microsatellite repeat is discarded. For example, in the sequence CGTGTCGTGT the two-copy repeat (CGTGT) is retained and the (GT) repeats are discarded. Microsatellite repeats can also overlap. For example, the sequence ACGACGACGTCGTCGT contains two microsatellite repeats, (ACG) and (CGT), which overlap by two characters, CG. In this case, the repeat boundaries are trimmed to avoid the overlap, but the repeat overhangs are not updated and the coded repeat character is kept the same. That is, the sequence ACGACGACGTCGTCGT will be represented by a repeat of type (ACG) coding for the sequence ACGACGA, then the non-repeating characters CG, followed by a repeat of type (CGT) coding for the sequence TCGTCGT. The overlap between two microsatellite repeats is always shorter than the shorter of the two motifs, and so trimming always leaves at least one nucleotide in the repeat and, therefore, cannot eliminate a microsatellite repeat. However, the final collapsed character may represent less than one copy of the repeat unit.

After graph construction, the reads are once again accessed to expand the microsatellite compressed strings by computing a consensus for each repeat. Reads are encoded as the original nucleotide string consisting of the nucleotides {A, C, T, G}, a stream of 16-bit integers representing the string in the microsatellite collapsed alphabet, and a character mapping that describes which substrings in the original string are represented by each character in the microsatellite collapsed string. The support for each encoded character is collected from all reads, and the most frequent nucleotide string is chosen as the consensus for each microsatellite collapsed character. If there is a tie, MBG selects the lexicographically highest one according to C++ string ordering. Note that after expansion of the microsatellites, the strings remain homopolymer compressed.

### Multiplex de Bruijn graph

MBG is a tool for building sparse de Bruijn graphs from long, accurate reads. Originally based on a minimizer ^62^ de Bruijn graph, the updated version of MBG uses closed syncmers ^63^ to sparsify the *k*-mer space and implements the multiplex de Bruijn graph strategy of Bankevich *et al*. ^15^ to simplify the graph. In summary, once the sparse de Bruijn graph is built for an initial size *k* (*k*=1001 by default), the reads are threaded through the graph according to exact *k*-mer matches. The threaded reads are then used to resolve the graph by locally increasing *k*. Bankevich *et al*. describe an exact de Bruijn graph with sequences stored on the edges. Here we describe how MBG adapts the multiplex de Bruijn graph algorithm to a sparse de Bruijn graph with sequences stored on the nodes.

Given a *k*-mer size *k*, all nodes with length *k* are potentially resolvable and taken under consideration. Paths which cross through a node are used to find spanning triplets for the potentially resolvable nodes. A spanning triplet for a node *n* is a subpath of length 3 where the middle node is *n*. The number of reads supporting each spanning triplet is found for all potentially resolvable nodes. Given a resolving coverage threshold *t*, if the read support for a spanning triplet is at least *t*, it is considered a solid triplet. By default a potentially resolvable node is marked unresolvable if any of its edges are not covered by a solid triplet, since an edge not covered by a solid triplet would be removed, introducing a gap in the graph. However, we disable this check for small *k* (empirically, *k* < 4,000) because these cases are usually caused by sequencing errors. Note that an edge is allowed to be covered by multiple solid triplets. After this, any solid triplets whose first and third nodes are of length *k* and marked unresolvable are removed. This process of marking nodes unresolvable and removing solid triplets is repeated until the set of potentially resolvable nodes and solid triplets has converged, and the final set of potentially resolvable nodes are marked resolvable.

Next, all resolvable nodes are removed and a set of edge-nodes, representing *k+1*-mers, is created. An edge-node is created for each edge touched by a resolvable node. If the edge touches one resolvable node, the edge-node has length *k+1*, containing the entire sequence of the resolvable node and one base pair from the successor node, and an edge is added between the newly created edge-node and the unresolvable node. If the edge connects two resolvable nodes, the sequence of the edge-node is the sequence of the path containing the two nodes. The edge-nodes are connected according to the solid triplets. Given a solid triplet (*n*_1_, *n*_2_, *n*_3_), an edge is added between the edge-nodes (*n*_1_, *n*_2_) and (*n*_2_, *n*_3_). The read paths are then rerouted to use these new edge-nodes, and non-branching paths are collapsed into a single node.

During the local *k* increase, MBG also performs graph cleaning. After each resolution step, tips and crosslinks are removed. A tip is a node with connections only on one end. Tips are removed if they are short (:s10 kb by default), low coverage (≥3 by default), and removing them would not create a new tip. Crosslinks are nodes which falsely connect two different genomic regions. We consider low coverage nodes with exactly one edge on each side as possible crosslink nodes and remove them if the tip removal condition applies to both ends of the node. After this, non-branching paths are again collapsed.

This process of increasing *k* and cleaning the graph is repeated until there are no more resolvable nodes or *k*=15,000, by default. MBG first resolves the graph with a coverage threshold *t*=2, and afterwards again with *t*=1 to resolve areas with lower coverage. Gaps in the graph caused by errors in the reads are then patched using the read paths, resulting in the final LA graph.

### Ultra-long read to graph alignment

After construction of the LA graph, we use GraphAligner^34^ to align UL reads to the graph for further resolution. Since the graph sequences remain homopolymer-compressed, all UL reads are compressed prior to alignment. During development, we noticed a low rate of systematic misalignments in GraphAligner and implemented changes which improved both accuracy and runtime compared with the previously published version.

First, to reduce memory requirements, we switched to maximal exact match seeding using an FM-index ^64^ instead of the default minimizer-based seeding. Second, the dynamic programming alignment formulation was systematically causing the alignment at the start of the read to be less accurate than the end of the read when aligning to bubbles in the graph. This was because the alignment was forced to pass through all the seeds, and when a seed was selected from the incorrect haplotype in a bubble, the previous implementation would not recover the optimal alignment. To address this, we adopted a Smith-Waterman formulation; however, to avoid computing the entire dynamic programming matrix, we implemented a sparse Smith-Waterman which restricts alignments from starting anywhere except for the start position of a seed match.

The sparse Smith-Waterman allows for the extension of seed hits from both sides of a bubble, without the runtime penalty of a full Smith-Waterman alignment. The runtime and memory requirements are further improved by banding. If there is a correct seed at the start of the read, the result of the sparse Smith-Waterman alignment is guaranteed to be optimal, regardless of the other seeds. The presence of correct seed hits near the beginning, but not at the start of the read will also have a high probability of recovering the optimal alignment. To counter the problem of spurious seeds at the beginning of a read, we also run the algorithm backwards, starting from the end of the read and extending towards the beginning, and the better of two alignments is taken.

In order to use a Smith-Waterman recurrence, the scoring matrix must have positive scores for good alignments and negative scores for bad alignments. However, GraphAligner uses Myers’ bit vector algorithm ^65^ which is based on edit distance, where positive scores correspond to bad alignments. We formulated a connection between edit distances and a scoring matrix suitable for Smith-Waterman. Given a desired identity threshold *p* and a dynamic programming matrix with the read represented by rows and the graph represented by columns, we define a match score *m* = 1, mismatch and deletion score *d* = -*p*/(1-*p*), and insertion score *i* = -*p*/(1-*p*) - 1 (with deletions and insertions defined relative to the graph sequence). Given this scoring scheme, an alignment consisting only of matches and mismatches with an identity of *p* will have an alignment score of 0, while higher identity alignments will have positive scores and lower identity negative scores. Given an alignment at row *n* in the dynamic programming matrix with an edit distance *e*, the edit distance can be used to compute the alignment score *s* = *n* - (*d*+1)*e* up to an additive factor. We use this equality to set the edit distance of alignment starting cells to a value which approximates an alignment score of 0. The identity threshold *p* also allows GraphAligner to discard alignments which are below the expected identity of highly accurate input reads. Alignments are also clipped based on alignment identity, something that cannot be easily done with an edit distance.

### Graph resolution with ultra-long reads

After the UL reads have been aligned to the LA graph, they are used to fill gaps and resolve repeats. The first step of the process is to connect nodes in the LA graph that were left disconnected due to coverage gaps or errors. Next, unique nodes are identified within the graph and connected based on the UL read paths. After this, the multiplex de Bruijn graph algorithm, identical to the one used by MBG, is run using the UL read paths to further resolve the graph.

When filling gaps in the LA graph, both the alignments of the UL reads and the topology of the graph are considered. First, tips are detected in the graph (i.e. nodes which don’t have an edge on both sides). If a UL read has an alignment ending at a tip, and another alignment starting at a tip, the UL read supports a gap fill between those two tips. If a pair of tips has a gap fill supported by a minimum number of UL reads (3 by default), a new gap node is inserted into the graph and connected to the tips. Each UL read supporting a gap fill has a gap length, estimated by the positions of the alignments within the read. Due to the higher error of the UL reads, there is no requirement that all gap fills have the same length. The length of the gap is estimated by taking the median gap length of all UL reads supporting the gap fill. If the gap has a positive length, the sequence of the gap node is represented by a corresponding number of “N”s. If it has a negative length, the sequences are copied from the tip nodes, and the edges are marked to have an appropriate overlap.

Unique nodes are those with a sequence that appears exactly once in the diploid genome. They are useful for identifying the correct genomic traversal of the graph, and Verkko uses multiple heuristics to discover unique nodes. The average genomic coverage is first estimated as the average LA read coverage observed for nodes ≥2100 kb. All nodes are then compared to this average coverage, and any node which is both long and close to the average coverage is marked as unique. Next, any node close to the average coverage, regardless of length, is marked as unique if it is path consistent. A node is considered path consistent if at least 80% of the UL read paths touching it are either identical, prefixes of each other, or suffixes of each other. The topology of the graph is then used to discover chains of superbubbles ^66^. Chains of superbubbles are classified as 1-copy, 2-copy, or multi-copy based on the coverage of the core nodes in the chain (i.e. the nodes separating the superbubbles). The core nodes of 1-copy chains are marked unique. For 2-copy chains, if a bubble has two paths with roughly equal coverage close to half the chain coverage, then the bubble nodes are marked unique. Multi-copy chains are ignored.

Once unique nodes are marked, the UL reads are used to find bridging paths between them. A bridge connects two unique nodes, with no unique nodes in between. The subpaths of UL alignments are collected as bridges and inconsistent bridges are resolved. Two bridges are considered inconsistent if they share one, but not both, unique node endpoints. At this stage, only the endpoints of the bridges are considered and not the specific paths. If an inconsistent bridge has less than half the read support of another, it is assumed to be erroneous and removed. If neither of the bridges has twice the coverage of the other, they are both retained. Once the bridges connecting the unique nodes are found, Verkko looks at the paths connecting the endpoints. The path with the most read support between each pair of unique nodes is taken as the consensus bridge path. All paths with at least half the coverage of the consensus path are kept, and all other lower coverage paths are discarded.

Lastly, tangles are marked as resolvable if all unique nodes touching the tangle have a bridge, and all high-coverage nodes and edges in the tangle are covered by a bridge (node coverage is measured from both the LA graph and aligned UL reads, and edge coverage from the aligned UL reads). These tangles are resolved by connecting the unique nodes based on the bridges. Nodes in a resolvable tangle that are not covered by a path are assumed to be erroneous and removed, while non-unique nodes covered by multiple paths are duplicated onto the corresponding paths. Unresolvable tangles for which there are no unique nodes, or those where the unique nodes were misidentified by the coverage checks, are left unchanged.

After the unique nodes have been connected, the multiplex de Bruijn graph algorithm, identical to the one implemented in MBG, is applied to the graph using the UL read paths. This can further resolve tangles which were not resolved by the unique node connection, either completely or partially.

### Graph cleaning

Once UL resolution is complete, tips are clipped from the graph. If there is at least one long path starting at a fork, then other shorter paths starting at the same fork may be removed. By default, long paths are ≥235 kb, and the clipped paths are either <35 kb or <10% of the longer path’s length, whichever is shorter.

Each step of resolution outputs a node mapping which relates the nodes in the previous graph to the simplified graph. This node mapping can be used recursively to discover which nodes in the original MBG graph are contained in a node at any point during resolution. This information is used to generate coverage estimates for each node in the subsequent graphs. The coverage estimate of a node is defined as the weighted average of the coverages of the MBG nodes contained in it, considering only MBG nodes contained in a single node in the final graph.

These coverage estimates are used for resolving bubbles caused by sequencing errors. If a chain of bubbles appears to be single-copy, any bubbles comprising it must be caused by sequencing errors. The highest coverage path through a bubble is kept and the nodes and edges outside of this path are discarded. The final graph combining both the LA and UL reads is named the ULA graph.

### Haplotype reconstruction

Rukki (“spinning wheel” in Finnish) is a companion tool for extracting haplotypes from a labeled assembly graph. Its main purpose is to facilitate haplotype-resolved assembly of diploid genomes by analysis of haplotype-specific markers within the graph nodes. For now, we have primarily targeted trio-based haplotype reconstruction, using parent-specific *k*-mers identified from parental Illumina reads as haplotype markers (Supplementary Note 1).

Rukki first labels graph nodes as maternal or paternal based on the prevalence of the corresponding haplotype markers, or leaves them unlabeled if the markers are ambiguous. This step considers an absolute number of markers attributed to the node, their average density, as well as the ratio between the marker counts of the two haplotypes. Rukki also detects nodes with a high abundance of markers from both haplotypes, as these nodes are likely to be misassembled or represent spurious cross-haplotype connections.

We call a graph node “solid” if its length exceeds a certain threshold (>500 kb). This threshold is empirically chosen so that solid nodes are common, but are very likely to be single-copy and haplotype-specific. Next, Rukki labels homozygous nodes by identifying neighborhoods where differently labeled regions converge. Due to minor flaws in the graph, nodes representing homozygous regions may contain a few parent-specific markers, which can lead to their initial mislabeling, so both unlabeled and previously labeled nodes with elevated coverage (as compared to the weighted average across all solid nodes) are examined. The remaining solid nodes are used to seed heuristic extension of the haplotype paths. Naturally, paternal nodes are considered incompatible with any path derived from a maternal node and vice versa, while homozygous and unlabeled nodes can potentially be incorporated in any path.

To make the extension procedure more robust to spurious cross-haplotype nodes, every time a solid node *s* is incorporated into the path, Rukki analyzes its neighborhood to try and identify the next solid candidate *t*. To do so, Rukki considers a subgraph bounded by solid nodes, with *s* being one of the sources. If all sources and sinks in the subgraph are labeled, and there exists only a single source and sink compatible with the current path’s haplotype (with *s* being the source), the corresponding sink is marked at the next solid candidate *t*. If found, Rukki further constrains the extension heuristic to prioritize nodes along any paths connecting *s* to *t*. Note that we only need to consider extensions in the forward direction, since a backward extension can be expressed via forward extension of the corresponding reverse complement path. The haplotype path extension can infer some missing haplotype labels, so the entire process is repeated a second time to generate a final set of haplotype paths. Supplementary Figure 23 provides some examples of haplotype paths produced by Rukki.

A unique feature of Rukki is its ability to “scaffold” across local ambiguities and breaks in the assembly graph, without the use of any additional scaffolding technologies. For example, when the extension procedure reaches a superbubble it will either completely traverse it (Supplementary Fig. 23A) or introduce a gap, depending on the bubble characteristics and provided options. Another example is scaffolding across a “loop tangle”, typically caused by near-identical tandem repeats and defined as a strongly connected subgraph with one source and one sink (Supplementary Fig. 23B). In this case the gap size is estimated based on the total length of the nodes in the tangle and their overlaps. Rukki can also scaffold across a coverage break in one haplotype when the other haplotype is intact (Supplementary Fig. 23C). In this case, the complete haplotype is used to estimate the size of the gap in the broken haplotype.

Rukki is an independent module taking as input only an assembly graph in GFA ^67^ format and the counts of haplotype-specific markers assigned to each node. Thus, it should be compatible with alternative methods of graph construction and haplotype assignment (e.g. based on Hi-C and Strand-seq). However, Rukki’s current heuristics have been optimized for Verkko’s ULA graphs, and would likely need further tuning for other graphs and contexts. Regarding alternative sources of phasing information, note that Rukki expects exactly two haplotype labels, which in the case of trios correspond to the paternal and maternal haplotypes. Fortunately, most phasing methods produce exactly two sets of contigs or read bins, typically labeled H1 and H2. The preliminary Hi-C integration results presented here are based on markers extracted from reads binned by DipAsm ^41^, but one could also use phased contigs from Hifiasm (Hi-C) or PGAS ^18^ (Strand-seq). A more sophisticated incorporation of phasing data is left as future work, and would likely involve mapping the Hi-C or Strand-seq reads directly to the Verkko assembly graph. Phased polyploid assembly is also left as future work and would require modifications to both Verkko and Rukki.

### Consensus

Verkko’s primary data structure is a homopolymer-compressed assembly graph so a consensus stage is required to recover the genomic sequence of individual nodes or haplotype paths. Verkko’s node mapping is used to translate each node or path in the final graph into a sequence of MBG nodes. The sequences of MBG nodes and the paths of the LA reads in the MBG graph are used to build a layout of reads for each node in the final graph. Each LA read is assigned to the location with the longest exact match through the MBG graph. If there are multiple equally good locations, the read is assigned to all of them. Due to their lower quality, UL reads are only included in the layout where they were used to fill a gap in the initial LA graph. A consensus sequence is then called for the reads in the layout using a module from Canu^68^. By default, the ends of the consensus sequence supported by only a single read are trimmed.

## Data availability

No new data was generated for this study. All assemblies generated in this paper are archived at zenodo https://doi.org/10.5281/zenodo.6618379 and we have provided convenience links to download both data and assemblies at https://github.com/marbl/verkko/blob/master/paper/README.md

## Code availability

Verkko code is available from: https://github.com/marbl/verkko and all code used for the paper is archived at zenodo under https://doi.org/10.5281/zenodo.6618379.

## Acknowledgements

This work was supported, in part, by the Intramural Research Program of the National Human Genome Research Institute, National Institutes of Health (MR, SN, AR, BPW, AMP, and SK) as well as grants from the U.S. National Institutes of Health (NIH HG010169 and HG002385 to EEE) and National Institute of General Medical Sciences (NIGMS 1F32GM134558 to GAL). EEE is an investigator of the Howard Hughes Medical Institute. This work utilized the computational resources of the NIH HPC Biowulf cluster (https://hpc.nih.gov).

## Author contributions

Methods and software development: MR, SN, BPW, SK. Analysis and validation: GAL, DP, AR, SK. Resource provision: EEE, AMP. Manuscript draft: MR, SN, AMP, SK. Figures: MR, SN, GAL, DP, AMP, SK. Editing: MR, SN, BPW, AMP, SK with the assistance of all authors. Supervision: EEE, AMP, SK. Conceptualization: MR, SN, AMP, SK.

