## Supplementary Notes + Figures for "Verkko: telomere-to-telomere assembly of diploid chromosomes"

### Supplementary Materials for Verkko: telomere-to-telomere assembly of diploid chromosomes

Supplementary Note 1: Genome assembly 2
Supplementary Note 2: Validation 4
Supplementary Tables (external file) 7
Supplementary Figures 8

### Supplementary Note 1: Genome assembly

We used Verkko v1.0 beta2, Flye v2.9-b1774, Hifiasm v0.16-r375, and LJA v0.2 with the commands:

```
hifiasm -l0 -o asm -t 32 hifi/*.fastq.gz
flye -t 32 --out-dir asm --nano[-hq|raw] ont/*.fastq.gz
lja -o asm -t 32 `ls ../hifi/*.fastq.gz |awk '{print "--reads "$1}'|tr '\n' ' '`
verkko --slurm -d verkko_hifi --hifi hifi/*.fastq.gz
verkko --slurm -d verkko --hifi hifi/*.fastq.gz --nano ont/*.fastq.gz
```

All assemblies ran on the NIH biowulf (https://hpc.nih.gov) cluster, with all assemblers except Flye using nodes from the normal partition. Flye ran on the largemem partition due to its high memory requirements (Supplementary Table 1).

Due to the high coverage, the *A. thaliana* dataset was downsampled to 80x HiFi and 100x ONT. For HG002 and *H. axyridis* we removed the `-l0` parameter from Hifiasm and added the `--diploid` parameter for LJA. For Flye we used `--nano-hq` for *A. thaliana* and `--nano-raw` for *H. axyridis* and HG002 as they had older ONT data (prior to Guppy 5). We added `--h1 *R1_001.fastq.gz --h2 *R2_001.fastq.gz` for HG002 Hifiasm + Hi-C assembly and `-1 pat.yak -2 mat.yak` for Hifiasm + trio assembly. The yak databases were downloaded from: https://s3-us-west-2.amazonaws.com/human-pangenomics/index.html?prefix=submissions/6040D518-FE32-4CEB-B55C-504A05E4D662--HG002_PARENTAL_YAKS/HG002_PARENTS_FULL/

We attempted to run `--nano-hq` on CHM13 but Flye ran out of memory on a 1.5 TB node so we used the `--nano-raw` results instead. For HG002 we used the HPGP downsampled dataset (35x HiFi, 60x ONT UL) for all assemblers and then ran Verkko using the full-coverage dataset (105x HiFi, 85x ONT UL).

We re-ran LJA on CHM13 and HG002 to get timing information. We found these assemblies were similar to the published LJA results so we re-used published LJA [15] assemblies of HG002 and CHM13. We also used previously published trio and Hi-C (60x Hi-C and 90x Hi-C)[36] for Hifiasm. We found the 60x and 90x Hi-C assemblies to be comparable (NG50 52.79 vs 52.50; switch 0.36% vs 0.22%; errors 40 vs 42) and so presented only the 60x assembly statistics in the manuscript.

We used available parents for HG002 and Merqury [45] to identify parent-specific markers in homopolymer-compressed space:

```
cd <path to maternal data> (repeat for paternal data and child data)
ls *fastq.gz > input.fofn
mkdir scripts
cat $MERQURY/build/count.sh |sed 's/count/compress count/g' > scripts/count.sh
cat $MERQURY/_submit_build.sh |awk '{if (match($0, "count.sh")) print
"script=`pwd`/scripts/count.sh"; else print $0}' | sed 's/16g/24g/g' > submit.sh
sh submit.sh 30 input.fofn maternal_compressed

# find hapmers
sh $MERQURY/trio/hapmers.sh maternal_compressed.k30.meryl
paternal_compressed.k30.meryl child_compressed.k30.meryl
```

```
# color asm
cat 5-untip/unitig-popped-unitig-normal-connected-tip.gfa |awk ‘{if (match($1,
“^S”)) { print “>”$2; print $3}}’ |fold -c >
unitig-popped-unitig-normal-connected-tip.fasta
sh $MERQURY/trio/hap_blob.sh maternal.k30.hapmer.meryl paternal.k30.hapmer.meryl
5-untip/unitig-popped-unitig-normal-connected-tip.fasta assembly
awk -v F=0.90 'BEGIN{print "node\tmat\tpat\tmat:pat\tcolor";}$1!="Assembly"{color
= "#AAAAAA"; if ($3+$4 > 100) { if ($3 > ($3+$4)*F) { color = "#FF8888"; } else if
($4 > ($3+$4)*F) { color = "#8888FF";} else { color = "#FFFF00"; } }; print $2
"\t" $3 "\t" $4 "\t" $3 ":" $4 "\t" color;}' < assembly.hapmers.count >
assembly.hapmers.colors.csv
```

These markers were used to color the Verkko graph nodes and report a ratio, for each unitig, of maternal and paternal markers. For Hi-C data, we used the available binned reads [11,41], https://s3-us-west-2.amazonaws.com/human-pangenomics/index.html?prefix=publications/HG002_BAKEOFF_2021/H2M/DFCI_SG.HG002.HiFi-ONT.binned/) to generate haplotype-specific markers and colored the Verkko graph as above. Note that unlike the trio markers, these may not be consistent between chromosomes. We then generated GFA files w/o sequences and injected coverage:

```
cat 5-untip/unitig-popped-unitig-normal-connected-tip.gfa |awk '{if (match($1,
"^S")) print $1"\t"$2"\t*\tLN:i:"length($3); else print $0}' > tmp.gfa
python inject_coverage.py --allow-absent
5-untip/unitig-popped-unitig-normal-connected-tip.hifi-coverage.csv tmp.gfa >
5-untip/unitig-popped-unitig-normal-connected-tip.noseq.gfa
```

These were input to Rukki with the commands:

```
cargo run -- trio -g unitig-popped-unitig-normal-connected-tip.noseq.gfa -m
assembly.hapmers.colors.csv --gaf-format -p out_paths.gaf -init-assign
out_init_ann.csv --refined-assign out_refined_ann.csv --final-assign
out_final_ann.csv --marker-sparsity 5000 --issue-sparsity 1000 --issue-len 200000
--try-fill-bubbles --marker-ratio 5. --issue-ratio 3. --issue-cnt 100
```

for trios and

```
cargo run -- trio -g unitig-popped-unitig-normal-connected-tip.noseq.gfa -m
assembly.hapmers.colors.csv --gaf-format -p out_paths.gaf -init-assign
out_init_ann.csv --refined-assign out_refined_ann.csv --final-assign
out_final_ann.csv --marker-sparsity 5000 --issue-sparsity 1000 --issue-len 200000
--try-fill-bubbles --marker-ratio 3. --issue-ratio 2. --issue-cnt 1000
```

for Hi-C. This produced scaffold paths. A new read layout and consensus was generated using Verkko:

```
verkko -d beta_asm --cnspath out_paths.gaf --cnsdir consensus
```

The Rukki step and consensus steps are now integrated into the Verkko pipeline. This required less than 300 CPU h and less than 160 Gb memory per dataset. For *Harmonia axyridis*, since no pre-binned Hi-C reads were available, we ran DipAsm commit 39d24cfd0ede139b209b4607c29c0ed2e6461fd9 with the command:

```
sh pipeline.sh hic ccs test FALSE out
```

With the ccs folder containing the HiFi reads and the hic folder containing the Hi-C libraries. We re-used the primary contigs of the reference assembly rather than running Peregrine [69].

#### Supplementary Note 2: Validation

We used QUAST from https://github.com/snurk/quast, with updates to Minimap2's v2.24 [70] alignment parameters. All assemblies were validated with the command:

```
quast.py -t 16 --skip-unaligned-mis-contigs -e --scaffold-gap-max-size 5000000
--no-snps --min-identity 98. -r $ref --min-alignment 10000 --extensive-mis-size
2000 --min-contig 50000 --large $reference assemblies/*.fasta
```

The references were GWHBDNP00000000.1, available at the China National Center for Bioinformation website [35] for *A. thaliana*, CHM13 v1.1 [14] for CHM13, and GCA_020881995.1 which includes completed chromosomes X & Y for HG002.

For *A. thaliana,* we identified reference regions with consistent differences in multiple assemblies (3/5) as putative errors and combined them with the errors identified in the reference by VerityMap [38] (see below). We also substituted the Verkko assembly combining HiFi and ONT data as the reference and compared the total unfiltered quast errors (Supplementary Table 2).

For CHM13v1.1, we used publicly available SV location and error locations (from https://github.com/marbl/CHM13-issues) to filter QUAST errors with the quast_sv_extractor.py script from https://github.com/skoren/helen [4] with the command:

```
python quast_sv_extractor.py -q contig_reports/all*<assembly>*.tsv -s variants.bed
-c rdna.bed -d v1.1_issues.mrg.bed -t empty
```

We used Mashmap [71] with the commands:

```
mashmap -r chm13.v1.1.fasta -q assembly.fasta -f one-to-one --pi 95 -s 10000
cat mashmap.out |awk '{if ($NF > 99 && $4-$3 > 500000) print
$1"\t"$6"\t"$2"\t"$7}' |sort |uniq > translation
cat translation|sort -k2,2|awk '{if ($3 > 15000000) print $0}' |awk -v LAST="" -v
S="" '{if (LAST != $2) { if (S > 0) print LAST"\t"C"\t"SUM/S*100"\t"TIG; SUM=0;
C=0; } LAST=$2; S=$NF; SUM+=$3; C+=1; TIG=$1} END {print
LAST"\t"C"\t"SUM/S*100"\t"TIG;}' |awk '{if ($2 == 1 && $(NF-1) > 95) print $0}'
|sort -k4 > chr_completness
cat translation|sort -k2,2|awk '{if ($3 > 15000000) print $0}' |awk -v LAST="" -v
S="" '{if (LAST != $2) { if (S > 0) print
LAST"\t"C"\t"SUM/S*100"\t"MAX/S*100"\t"TIG; SUM=0; MAX=0; C=0; } LAST=$2; S=$NF;
SUM+=$3; if (MAX < $3) MAX=$3; C+=1; TIG=$1} END {print
LAST"\t"C"\t"SUM/S*100"\t"MAX/S*100"\t"TIG;}'  |awk '{print $1"\t"$4}' |sort
-nk1,1 -s > chr_completeness_max
```

to map assemblies, select large alignments, and evaluate coverage of each chromosome by the largest unitig and overall completeness. We used the VGP [10] telomere pipeline and BEDTools [72] to identify canonical vertebrate telomeres with the commands:

```
find assembly.fasta > tmp.telomere
java FindTelomereWindows tmp.telomere 99.9  |awk '{if ($NF > 0.5) print
$2"\t"$4"\t"$5"\t"$3"\t"$NF}'|sed s/\>//g|bedtools merge -d -500 -i - -c 4 -o
distinct  |bedtools sort -i -|bedtools merge -i - > telomere
```

We used BACs available for CHM13 to validate the assemblies. Three known bad BACs [14] were excluded and validation was using bacValidation (https://github.com/skoren/bacValidation) with the command:

```
sh getStats.sh assembly.fasta
```

Reference-free validation was performed via VerityMap git commit 30673e8f4e10ce21ad653eec03a969d58593194a using the HiFi data available for each dataset with the command:

```
veritymap --reads hifi.reads.fastq -d hifi -t 32 -o output_asm
```

For *A. thaliana*, the high coverage allowed an independent set of HiFI reads to be used for validation that were not used for assembly. We reported errors by counting entries in the <asm>_kmers_dist_diff.bed when the allele frequency was at least 25 and the length was over 2 kb (to match QUAST parameters). Note that this may include some false-positive heterozygous variants. We could not use VerityMap on the full HG002 assemblies as it does not yet support alignments to diploid human genomes due to a lack of unique *k*-mers. Instead, we trio-binned all available HiFi data and randomly sorted the unassigned reads to maternal and paternal bins. The assemblies were separated using markers into maternal and paternal haplotypes and each was validated separately using the appropriate bin of reads.

We used Merqury [45] to evaluate switch errors using trio information on HG002 and QV on HG002 with a k=21 Illumina database with the commands:

```
sh $MERQURY/trio/hap_blob.sh haplotype-MOM.meryl haplotype-DAD.meryl
assembly.fasta assembly
sh $MERQURY/trio/hamming_error.sh assembly.hapmers.count assembly.fasta
sh $MERQURY/eval/qv.sh HG002.k21.meryl assembly.fasta
```

We combined both haplotypes when running the above commands. To measure the frequency of parental markers in centromeres versus other regions of the genome, we looked at the count of markers versus positions chromosome-wide versus within the centromeric region. In the chromosomes overall 1.3% of the positions intersected a parental marker. Within the centromere, an average of 15% of  the positions intersected a parental marker. Due to incomplete UL integration, Verkko often outputs short unitigs with low quality. Thus, any unitig less than 1 Mb were excluded when computing QV.

We used asmgene (https://lh3.github.io/2020/12/25/evaluating-assembly-quality-with-asmgene) with the cDNA downloaded from embl (http://ftp.ensembl.org/pub/release-105/fasta/homo_sapiens/cdna/Homo_sapiens.GRCh38.cdna.all.fa.gz) and the CHM13 v1.1 assembly as the reference[14] with the commands:

```
minimap2 -cxsplice:hq -t$cores chm13.v1.1.fasta Homo_sapiens.GRCh38.cdna.all.fa >
ref.cdna.paf
minimap2 -cxsplice:hq -t$cores $asm Homo_sapiens.GRCh38.cdna.all.fa >
$prefix.cdna.paf
paftools.js asmgene -i0.97 -a ref.cdna.paf $prefix.cdna.paf > $prefix.stats
```

The missing duplicated from asm, complete single-copy also single-copy in asm, and complete single-copy not single-copy in asm were then computed as:

```
missing_duplicated=`cat $prefix.stats|grep full_sgl|awk '{print 100*$4/$3}'`
missing_single_copy=`cat $prefix.stats |grep dup_cnt|awk '{print
100*(1-($4/$3))}'`
duplicated_single_copy=`cat $prefix.stats |grep full_[sd] | awk '{if (match($0,
"full_sgl")) {R=$4; } else print 100*$NF/R}'`
```

Previously published Strand-seq data for HG002 [48] were aligned separately to paternal and maternal Verkko assembly of HG002 using BWA [73] aligner (version 0.7.17-r1188) and SAMtools [74] (version 1.10). We note that in order to alleviate spurious read mappings in pseudoautosomal regions of chromosome X and Y we have added assembled chromosome Y to maternal assembly and chromosome X to paternal assembly. Duplicate reads were marked using sambamba (version 1.0). Aligned BAM files were processed using breakpointR [75] in order to create haplotype specific Strand-seq composite files as previously described [48]. We again run breakpointR on such composite files to detect changes in read directionality with the following parameters: `windowsize = 50000, binMethod = "size", pairedEndReads = FALSE, genoT = 'binom', background = 0.1, peakTh = 0.25, minReads = 50`. Misorientation and unresolved homozygous inversions are reported as regions with the majority of minus oriented reads ('ww', Watson-Watson strand state) as one would expect all Strand-seq reads for a given sample to map in plus orientation ('cc', Crick-Crick strand state) if the assembly is correctly oriented. Regions genotyped as a mixture of Watson and Crick reads ('wc' - Watson-Crick strand state) are either caused by low mappability regions or regions where short Strand-seq data are not able to distinguish between paralogous sequence copies. Number of such regions represent true heterozygous inversion between paternal and maternal haplotype. To come to a final number of Strand-seq predicted heterozygous inversions we have selected regions with Watson reads fraction between 0.4 - 0.6 and regions with a predicted normal copy number.
Another level of assembly validation relies on Strand-seq ability to assign assembled contig or scaffolds to a chromosome of origin based on coinheritance of Strand-seq read directionality across multiple Strand-seq libraries [17,18]. This analysis was done by running SaaRclust [18] function 'scaffoldDenovoAssembly' with the following parameters: `pairedEndReads = TRUE, bin.size = 200000, step.size = 200000, prob.th=0.25, min.mapq = 10, bin.method = 'dynamic', min.contig.size = 100000, num.clusters = 100, ord.method = 'greedy', remove.always.WC = TRUE, eval.ploidy = TRUE, desired.num.clusters = 24, max.cluster.length.mbp = 260`. With this analysis we are able to assign each contig to a single cluster that represents a single chromosome. Also one is able to detect chimeric contigs or scaffolds that wrongly connect distal parts of the same or different chromosomes.

### Supplementary Tables (external file)

**Supplementary Table 1.** Assembler runtime and peak memory usage. Time and peak memory usage was calculated using SLURM's sacct utility. For Verkko, the runtime was the sum of all jobs run on the cluster. The majority of Verkko's runtime is spent on ONT alignments (93 CPU h or 58% on *A. thaliana*, 3748 CPU h or 96% on *H. axyridis,* 6800 CPU h or 64% on CHM13, and 15,000 CPU h or 81% on HG002). We expect future improvements to GraphAligner will significantly reduce this runtime.

**Supplementary Table 2.** Verkko assembly of A. thaliana is more continuous and accurate than either ONT-only or HiFi-only assemblies. The NG50 is measured against the published reference size and reports the longest unitig which is needed to cover half the genome. Completeness is measured from QUAST as are mismatches/indels and structural errors after filtering for likely errors in the reference (Supplementary Note 2). >99.5% complete chromosome reports unitigs which have alignments to a single reference chromosome and cover at least 99.5% of the reference chromosome length. VerityMap errors are results using an independent set of HiFi reads to identify errors in the assembly (Supplementary Note 2).

**Supplementary Table 3.** QUAST unfiltered errors versus Verkko assembly or versus the published reference. All assemblers have fewer errors versus the Verkko version than the published reference, confirming it has better support.

**Supplementary Table 4.** HG002 assembly results using standard HiFi data. When compared to DeepConsensus results, Verkko's switch rate is reduced 3-fold while continuity improves. We found that the higher switch rate in Verkko in this case was primarily due to a single region on Chromosome 10 where a haplotype coverage gap caused a switch during ONT-based resolution.

**Supplementary Table 5.** Asmgene analysis on the haplotype-resolved assemblies. For Verkko, we took scaffolds marked as haplotype 1 or haplotype 2 and any unclassified unitigs ≥100 kb. We did not evaluate the assemblies which were not separated into haplotypes (e.g. Hifiasm, LJA, or Verkko unitigs) because all the genes would appear duplicated. Multicopy genes missed indicate incompletely assembled repeats. Both Hifiasm and Verkko have similar low rates of missed genes. Completeness metrics report the fraction of complete and complete but duplicated genes. Duplicated genes do not count in the complete percentage so false-duplications necessarily drop the complete percentage. The false duplications present within a haplotype for Verkko, especially in the case of Hi-C data, therefore lower the completeness metric.

**Supplementary Table 6.** Strand-seq identified heterozygous regions in both haplotypes. The list of regions is plotted in Supplementary Figure 16.

### Supplementary Figures

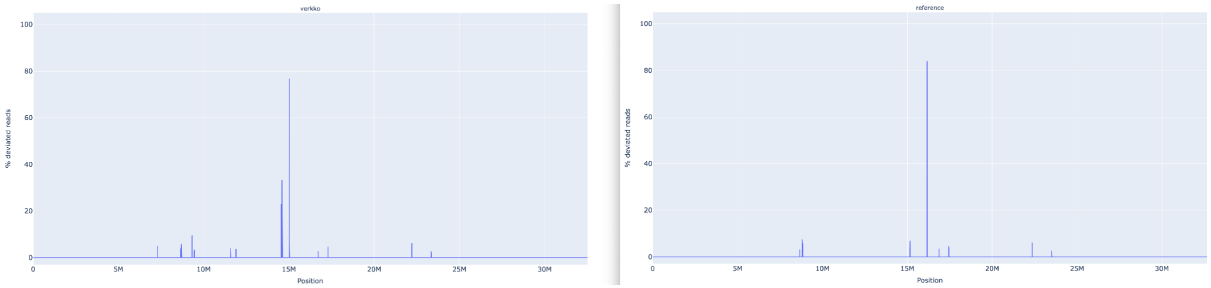


**Supplementary Figure 1.** *A. thaliana* Chr1 unitig in Verkko (left) vs published Chr1 evaluated by VerityMap [38]. VerityMap compares the spacing of unique *k*-mers within the HiFi reads to the spacing observed in the assembly. Whenever there is a disagreement, the plot shows a spike at the discrepant location. The x-axis indicates the coordinates along the assembly contig or scaffold while the y-axis shows the fraction of disagreeing reads (0-100%). A disagreement greater than 50% is likely not a heterozygous variant but a true error in the assembly. The BED file produced by VerityMap also indicates the size of the discrepancy, estimated from the difference in *k*-mer spacing between the reads and the assembly.

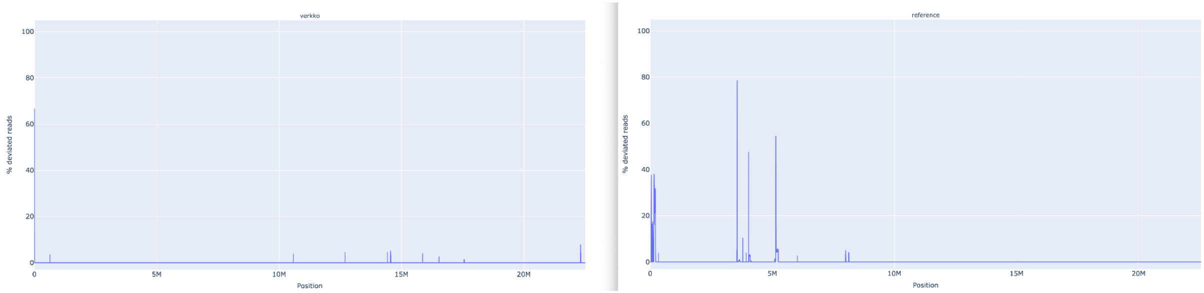


**Supplementary Figure 2.** *A. thaliana* Chr2 unitig in Verkko (left) vs published Chr1 evaluated by VerityMap.

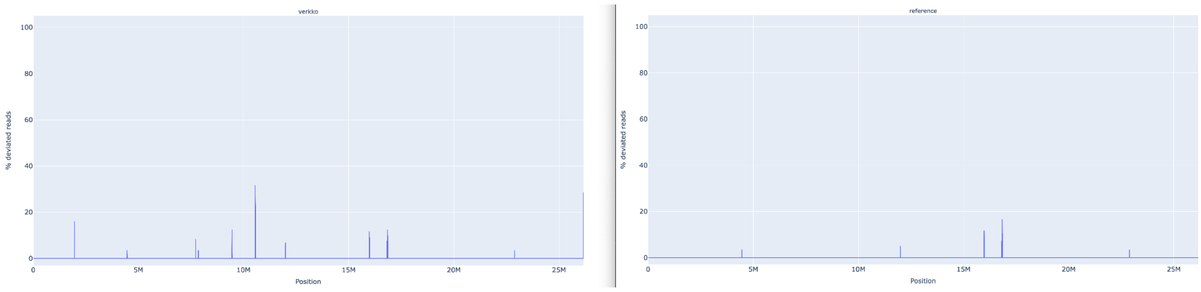


**Supplementary Figure 3.** *A. thaliana* Chr3 unitig in Verkko (left) vs published Chr1 evaluated by VerityMap.

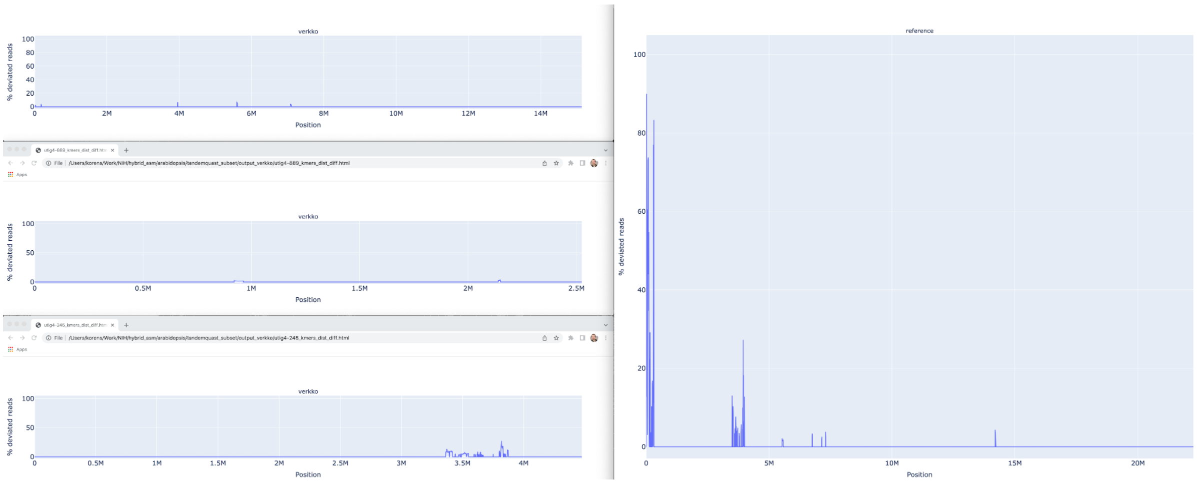


**Supplementary Figure 4.** *A. thaliana* Chr4 unitig in Verkko (left) vs published Chr1 evaluated by VerityMap.

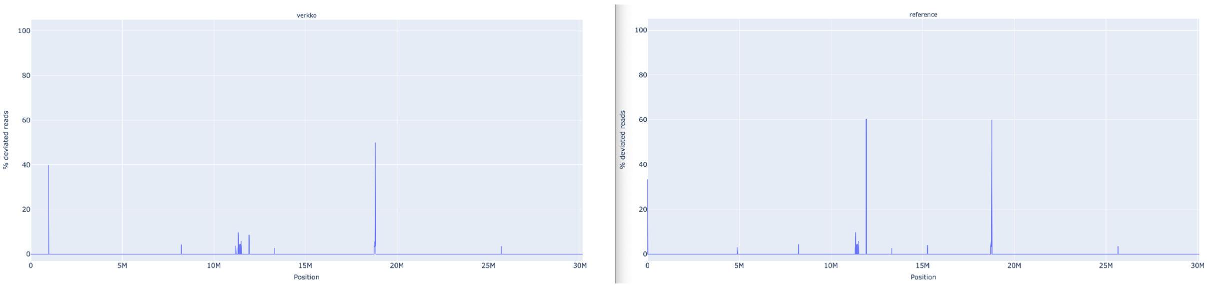


**Supplementary Figure 5.** *A. thaliana* Chr5 unitig in Verkko (left) vs published Chr1 evaluated by VerityMap.

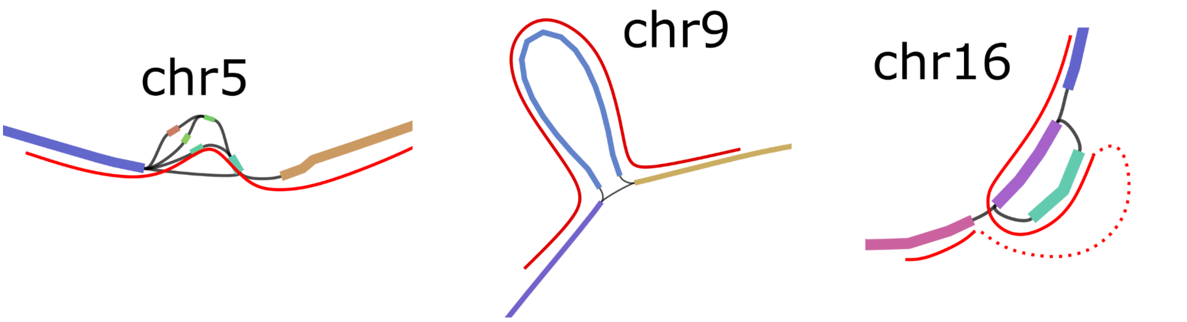


**Supplementary Figure 6.** The remaining unresolved regions in CHM13 chromosomes 5, 9 and 16, with the correct resolution marked in red paths. Left: Chr5 has a spurious edge causing a cycle, and three spurious low-coverage nodes which were not removed by bubble popping since they are a part of the cycle. Middle: Chr9 has a spurious edge. Right: Chr16 has two spurious edges, and one missing edge (dashed red curve).

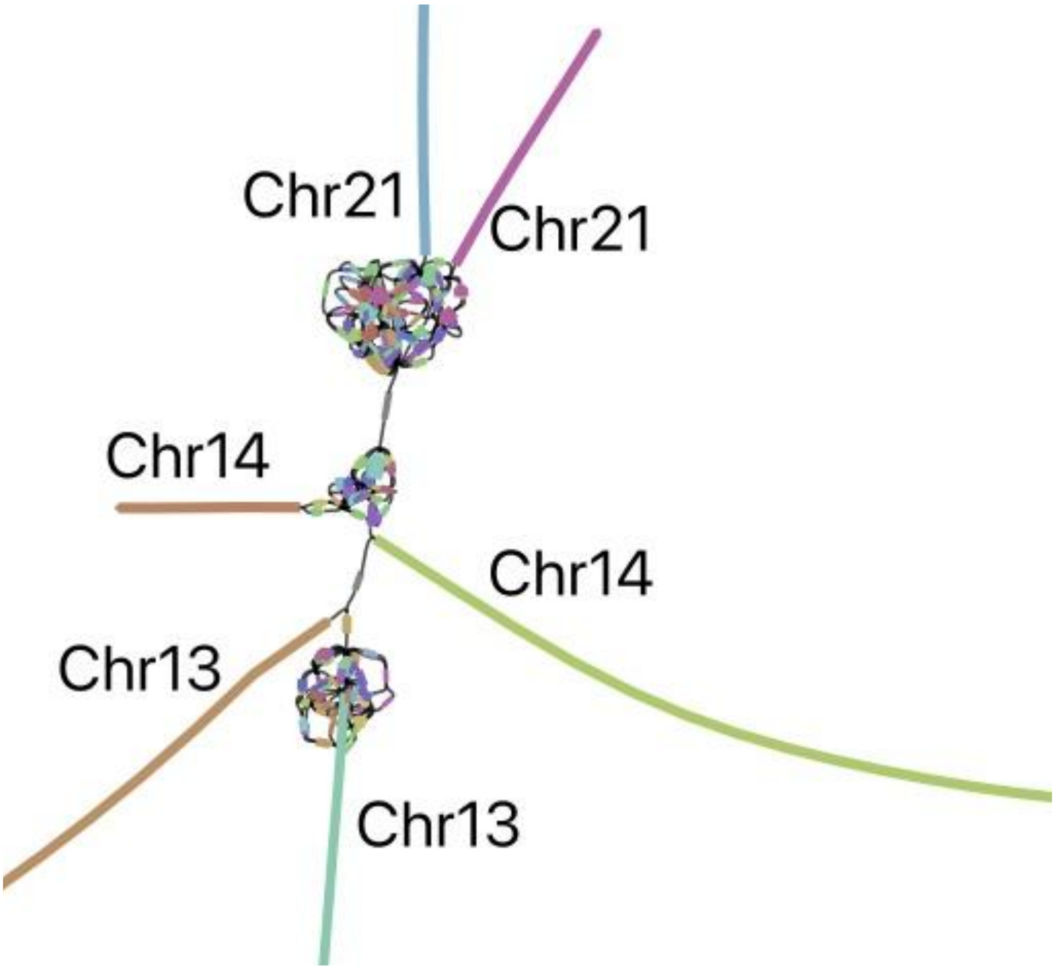


**Supplementary Figure 7.** rDNA cluster mixing in CHM13 chromosomes 13, 14, and 21. Each chromosome has a separate rDNA tangle. There are two cross-chromosomal connections by erroneous low coverage (<4x) nodes shown in gray. For all three chromosomes, the remainder of the p and q arms are contained in the long unitigs shown.

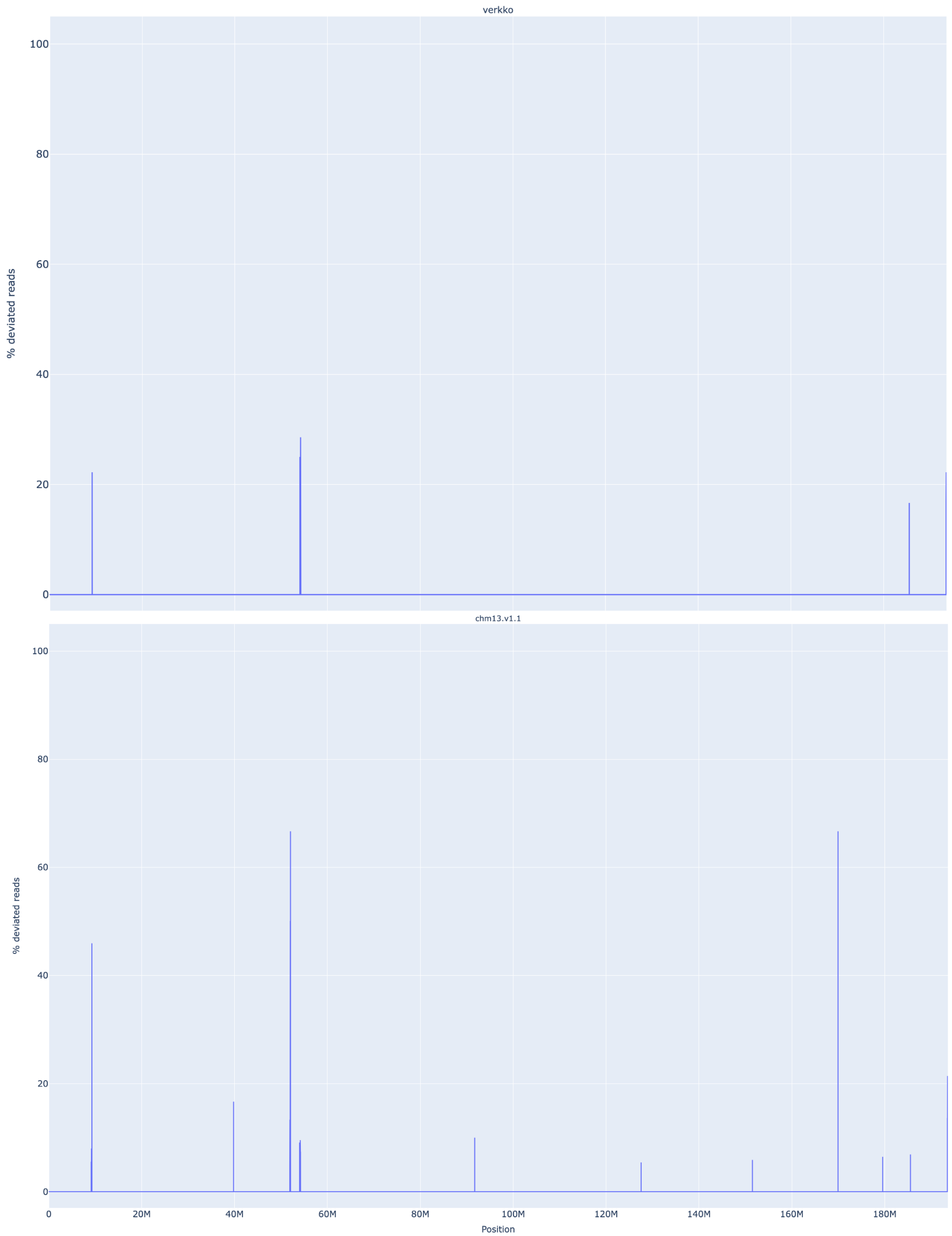


**Supplementary Figure 8.** VerityMap discrepant reads plot for CHM13 HiFi and ONT Chromosome 4 unitig assembled by Verkko (top) and CHM13 v1.1 [14] (bottom). The Verkko assembly has no regions where a large fraction of reads are deviated even though QUAST marks an error at approximately 52 Mb. This corresponds to a position in the reference with a large fraction of deviated reads and an estimated 19 kb discrepancy.

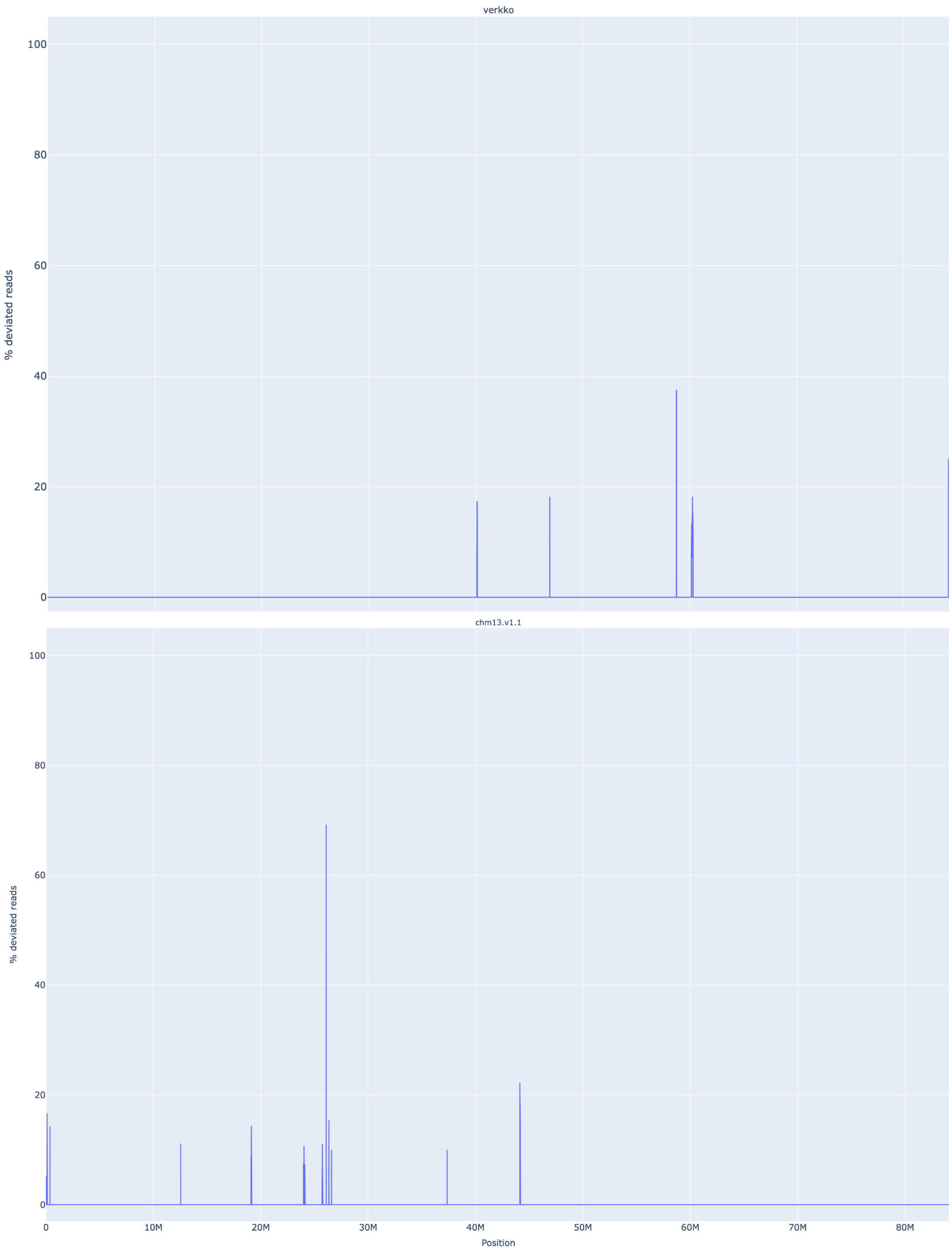


**Supplementary Figure 9.** VerityMap discrepant reads plot for CHM13 HiFi and ONT Chromosome 17 unitig assembled by Verkko (top) and CHM13 v1.1 (bottom). There are no regions with a large fraction (>50%) of discrepant reads in the Verkko assembly despite QUAST reporting an error at approximately 25 Mb on the reference. This corresponds to an approximately 3 kb discrepancy identified by VerityMap in CHM13 v1.1.

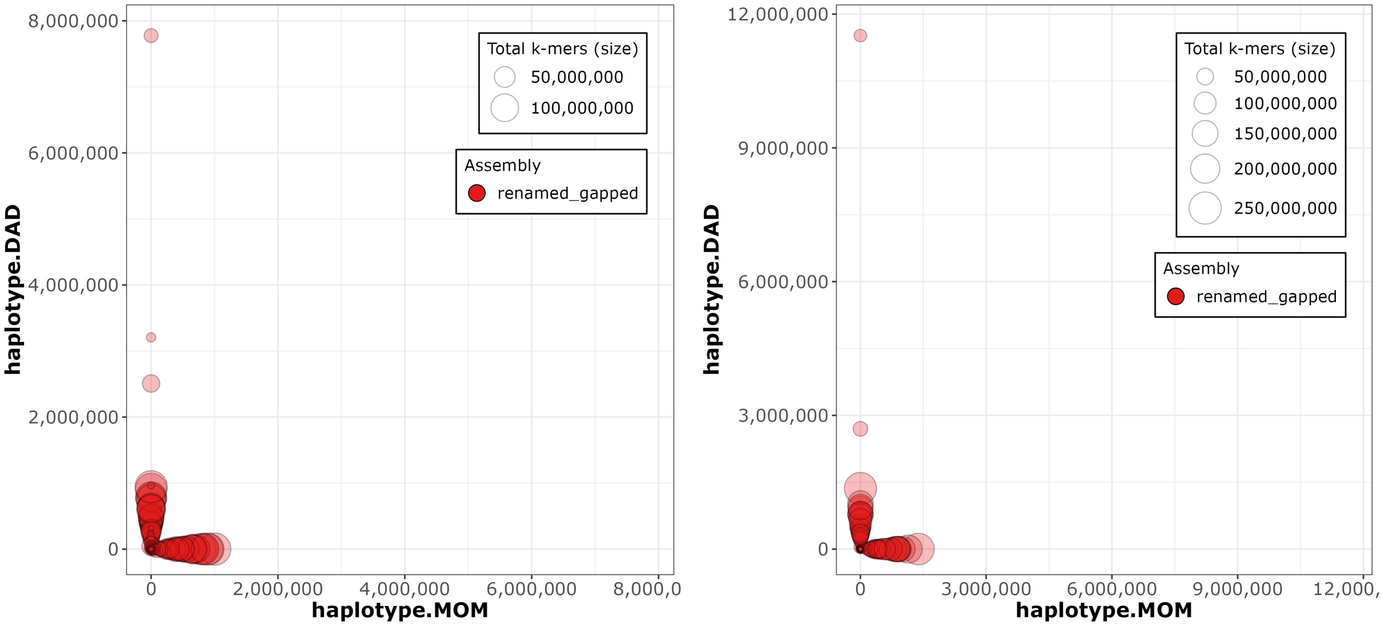


**Supplementary Figure 10.** Merqury [45] haplotype blob plots for HG002 downsampled Verkko Hi-C (left) and trio (right) assemblies. Each contig/scaffold is a circle on the plot, with the size scaled based on contig/scaffold length. The x-axis shows the number of maternal markers while the y-axis shows the number of paternal markers. Contigs which lie along either the x-axis or y-axis show no haplotype errors and are consistently maternal or paternal. Contigs which mixed haplotypes would appear along the diagonal but are not observed in these plots, indicating an accurately phased assembly.

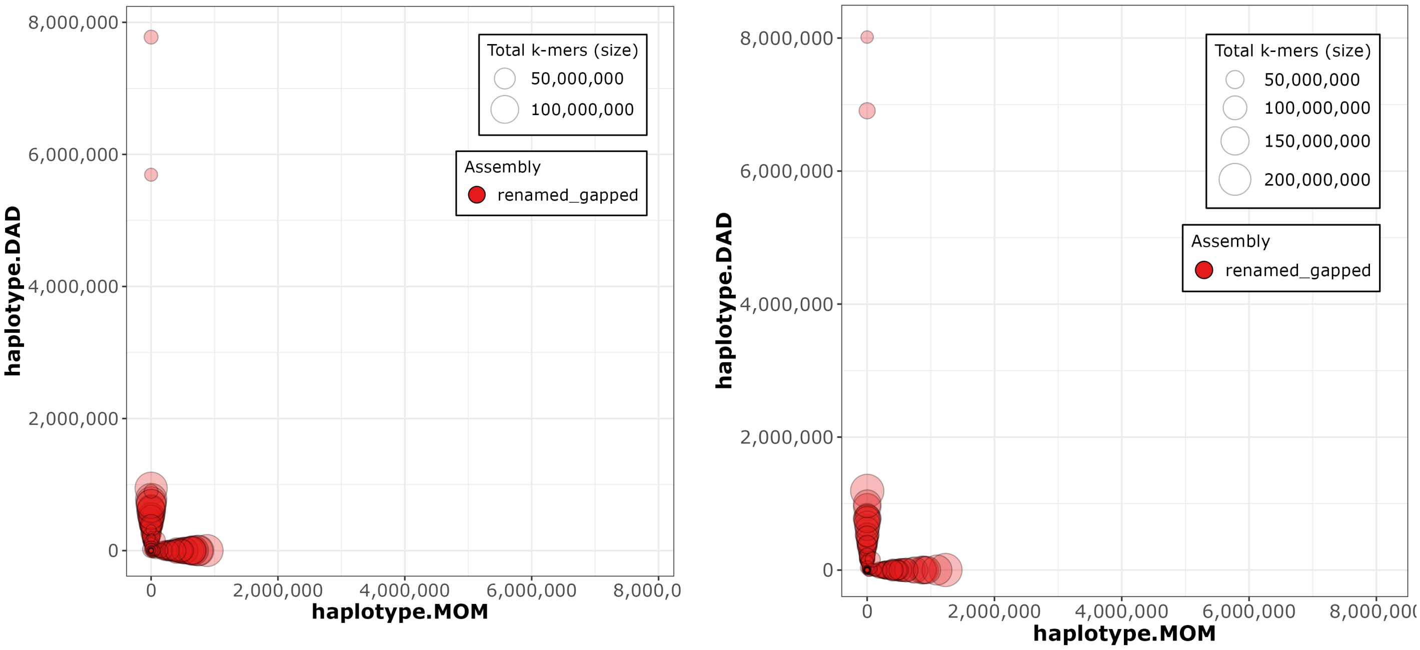


**Supplementary Figure 11.** Merqury haplotype blob plots for HG002 downsampled DeepConsensus HiFi Verkko Hi-C (left) and trio (right) assemblies.

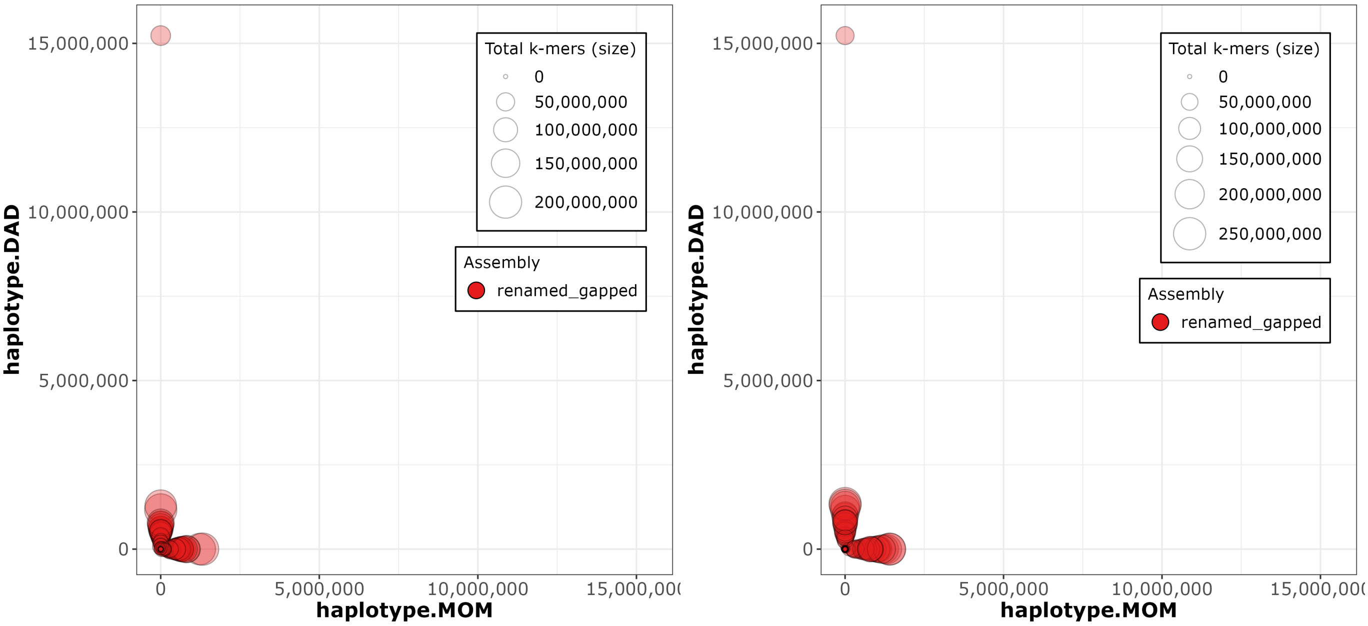


**Supplementary Figure 12.** Merqury haplotype blob plots for HG002 full-coverage Verkko Hi-C (left) and trio (right) assemblies.

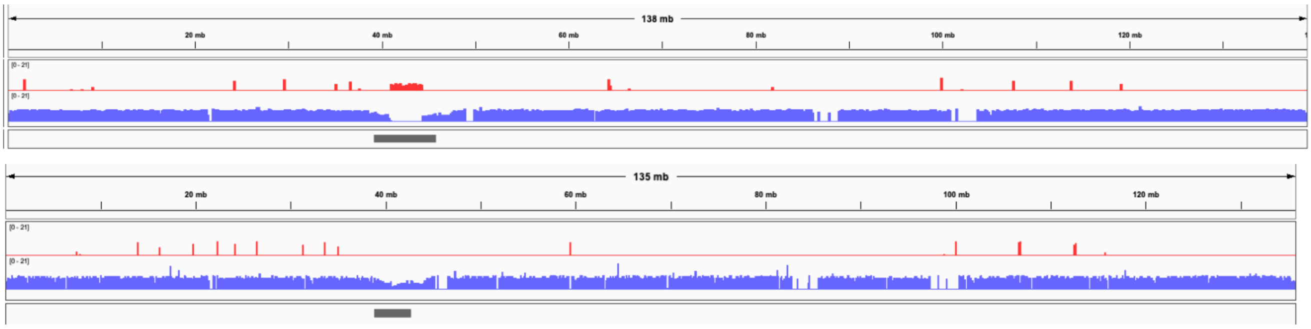


**Supplementary Figure 13.** IGV [76] views of a recently published HG002 diploid assembly of paternal Chromosome 10 [11] (top) and the Verkko full-coverage trio assembly of the same chromosome (bottom). The tracks show the maternal (red) and paternal (blue) markers. The centromere location is shown in gray. The published assembly has extensive switching within the centromere array, indicated by the presence of maternal markers and the absence of paternal markers. In contrast, the Verkko assembly centromere shows only paternal markers. The Verkko paternal centromere array is shorter but shows no signs of mis-assembly (Supplementary Fig. 19b) indicating the larger array in the published assembly is likely due to the incorrect insertion of maternal sequence. Overall, the Verkko assembly is more continuous, with 0 gaps vs 4, and a lower hamming error rate, 0.03%, versus 1.98% compared to the published assembly.

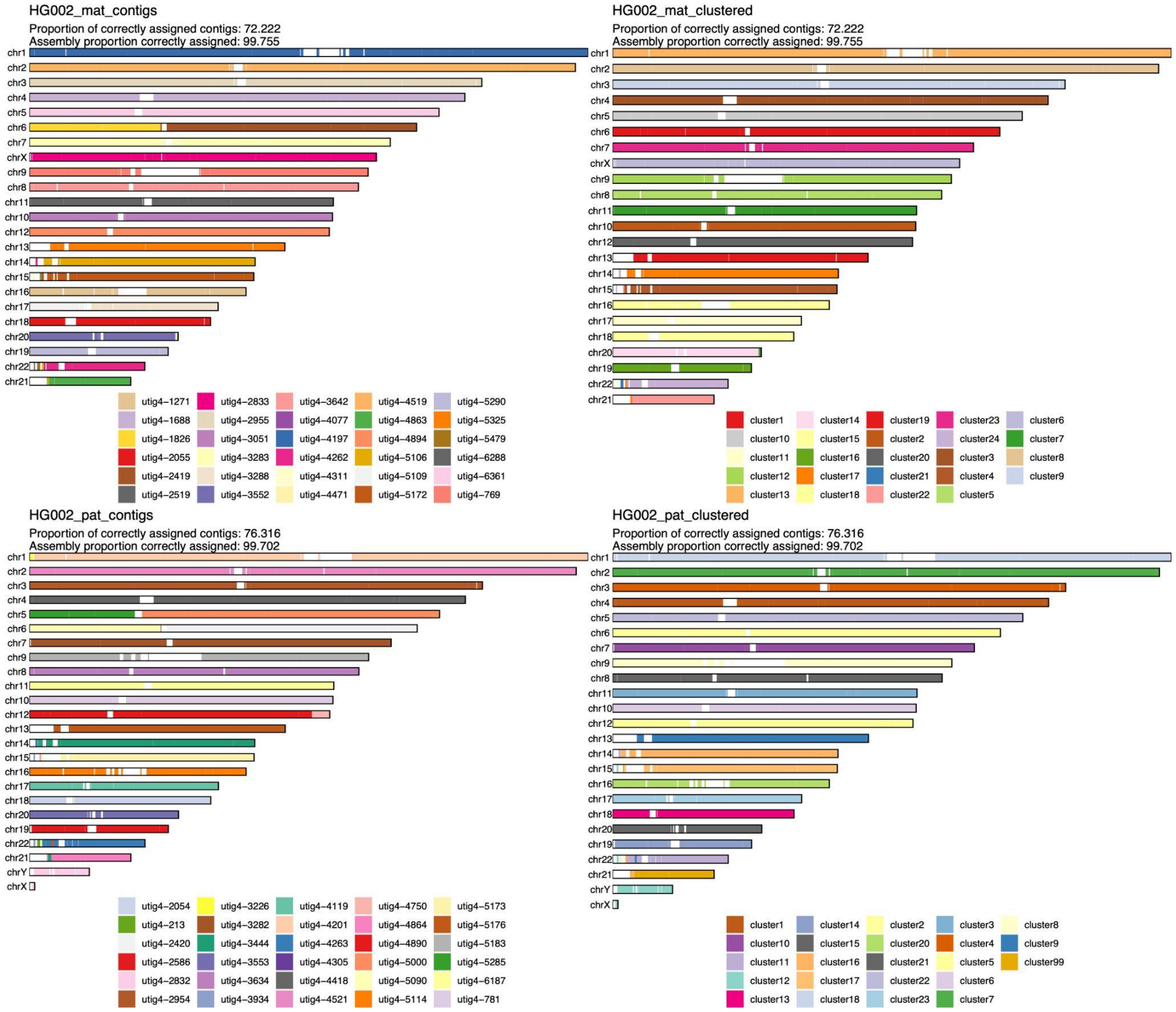


**Supplementary Figure 14.** Strand-seq validation of the full-coverage Verkko trio assembly. **Left:** alignment-based scaffold assignment to the maternal haplotype (top) and paternal haplotype (bottom). Almost all chromosomes are a single color, indicating that Verkko scaffolds resolved most chromosomes end-to-end. The only exceptions are in the acrocentrics, where some of the scaffolds could not be assigned due to low mappability and maternal Chromosome 6 and paternal Chromosomes 5 which are each composed of two large scaffolds. Over 99.7% of the scaffold bases could be assigned to chromosomes. **Right:** the cluster assignment for the maternal haplotype (top) and paternal haplotype (bottom) based on Strand-seq data. Here, cluster ID is assigned to each 200 kb window in a scaffold. In case of large scale chromosomal mis-joins, we expect to see multiple colors in a chromosome. The Verkko assembly is consistent with scaffolds all representing a single chromosome bin. Once again, >99.7% of the scaffold bases can be assigned using Strand-seq. Only 2 and 4 Mb of sequence not scaffolded by Verkko could be assigned to the maternal and paternal haplotypes, respectively.

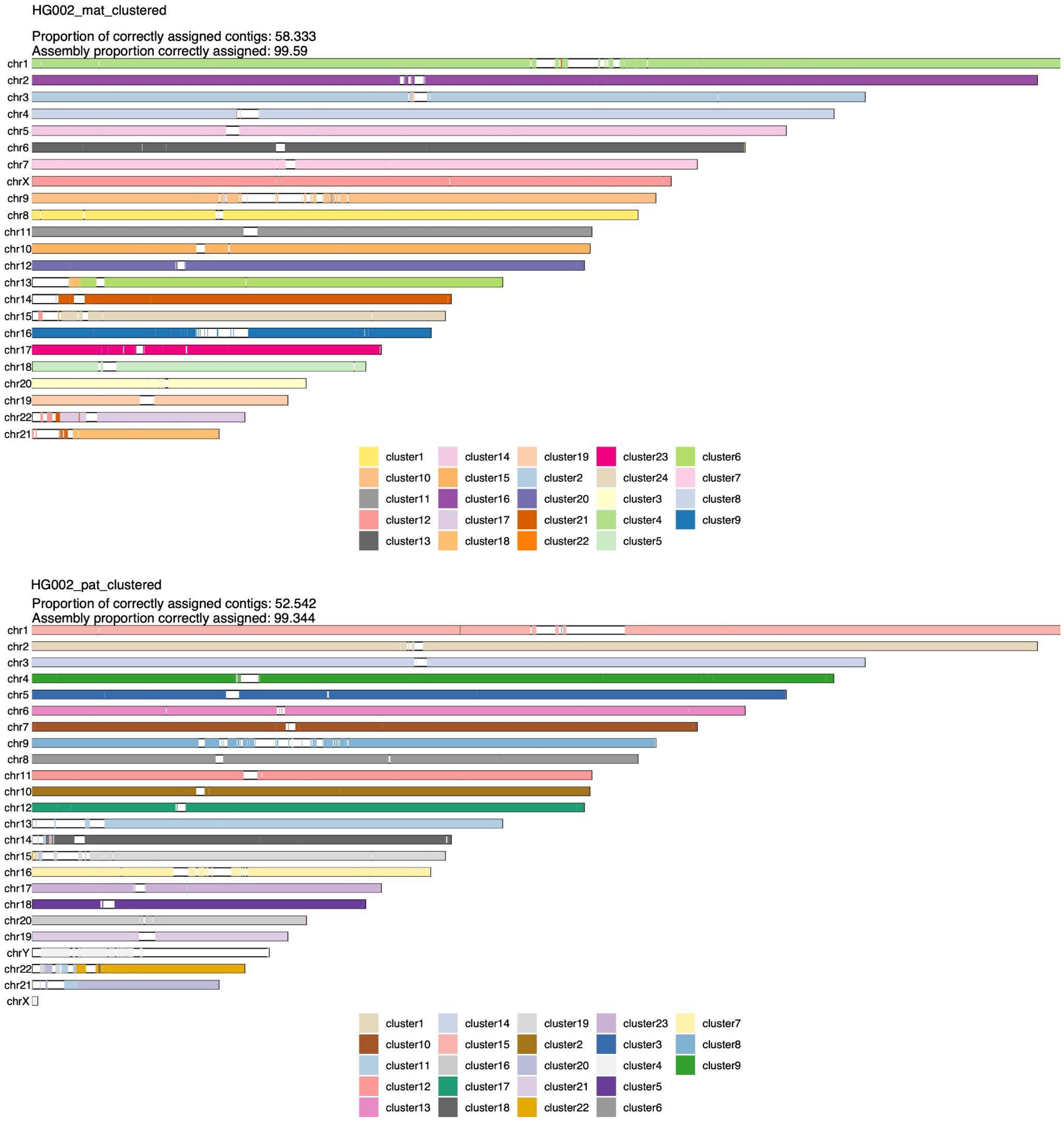


**Supplementary Figure 15.** Strand-seq validation of HPRC manually curated assembly [11]. The cluster assignment for the maternal haplotype (top) and the paternal haplotype (bottom) based on Strand-seq data. Here, cluster ID is assigned to each 200 kb window in a scaffold. In case of large scale chromosomal mis-joins, we expect to see multiple colors in a chromosome. A smaller fraction of contigs (and a slightly lower fraction of bases) was assigned than for the Verkko assembly in Supplementary Figure 14, despite the combination of technologies and manual curation. This may be due to shorter contigs from unresolved repeats which are resolved through Verkko's ONT integration. There is also visible chromosome mixing within the acrocentric chromosomes unlike in the Verkko result.

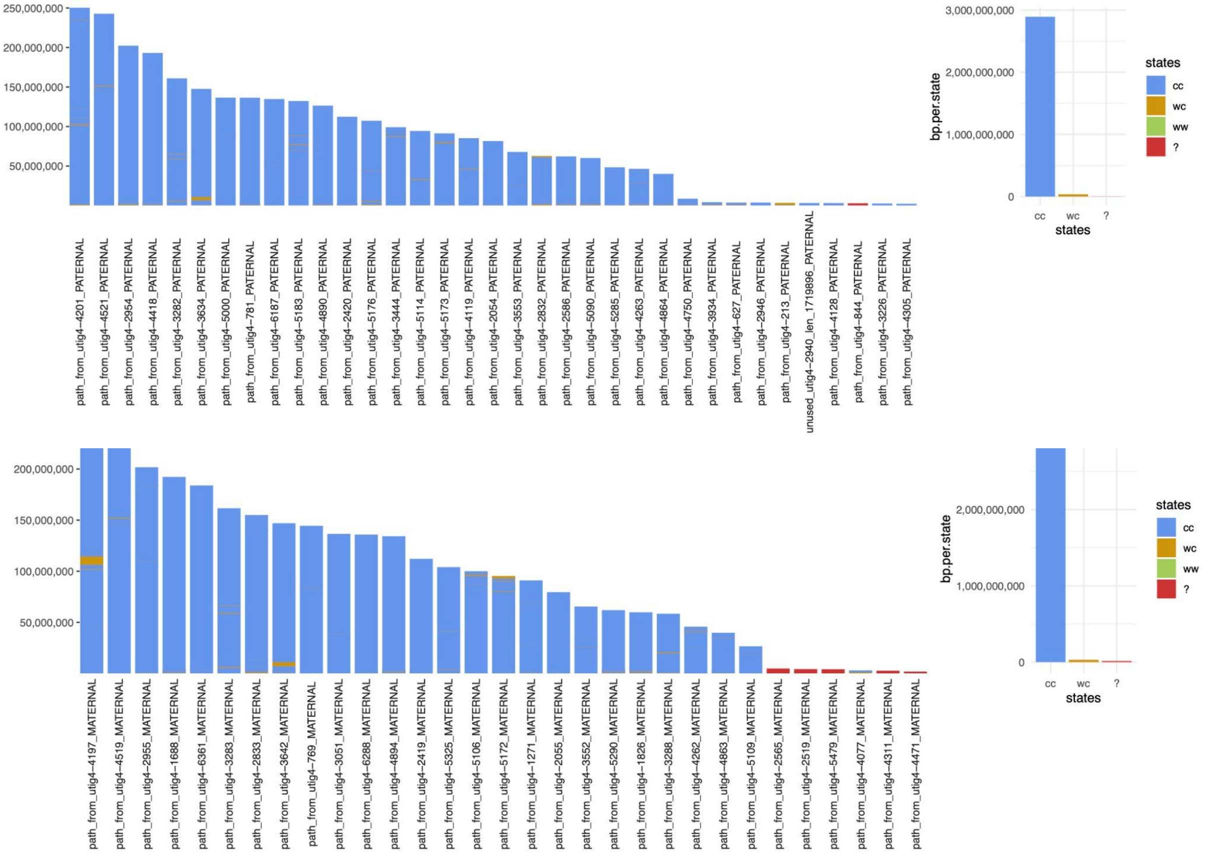


**Supplementary Figure 16.** The states assigned to each scaffold in the paternal (top) and maternal (bottom) for the full-coverage Verkko trio assembly. Strand-seq reads aligned to each assembly are genotype based on their directionality into three possible strand states. Crick-Crick ('cc') state in which both homologs in Strand-seq data map in direct orientation and thus such regions are consistent with Strand-seq directional information. Watson-Watson ('ww') state in which both homologs in Strand-seq data map in inverted orientation and are indicative of assembly misorientation or unresolved homozygous inversion. Lastly, there are a few (<1% of bases) Watson-Crick ('wc') where there is a mixture of Watson and Crick reads and such regions are indicative of heterozygous inversions between haplotypes or might also be cause by low-mappability regions for short Strand-seq reads.

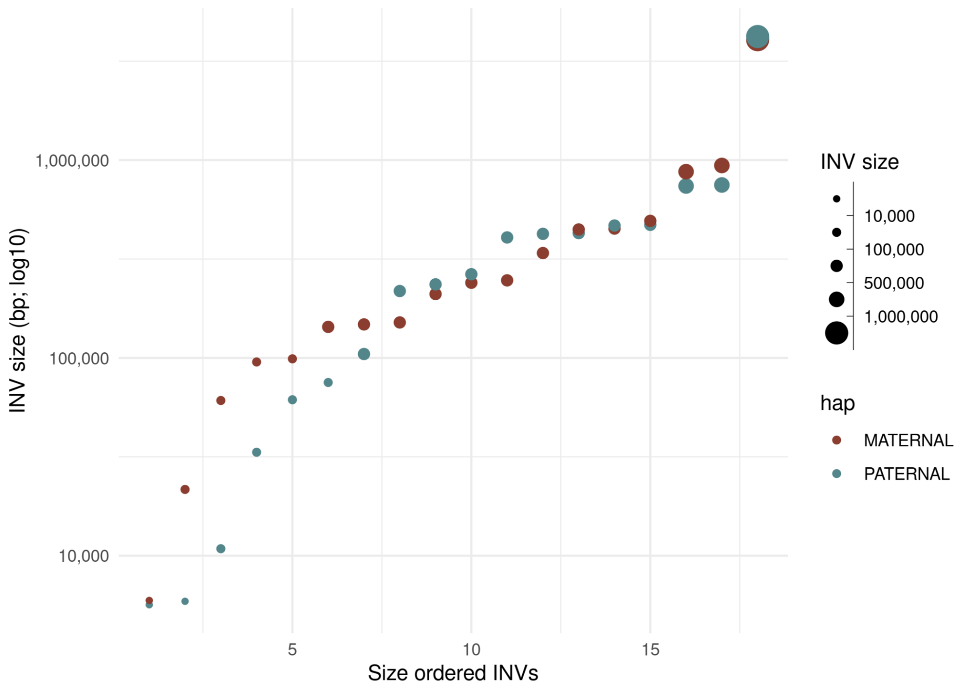


**Supplementary Figure 17.** The size of the heterozygous inversion versus the count of inversions of that size in the maternal and paternal haplotypes of the full-coverage Verkko trio assembly. These regions have confident Strand-seq alignments and normal copy number so these regions indicate potential true heterozygous variation between the haplotypes.

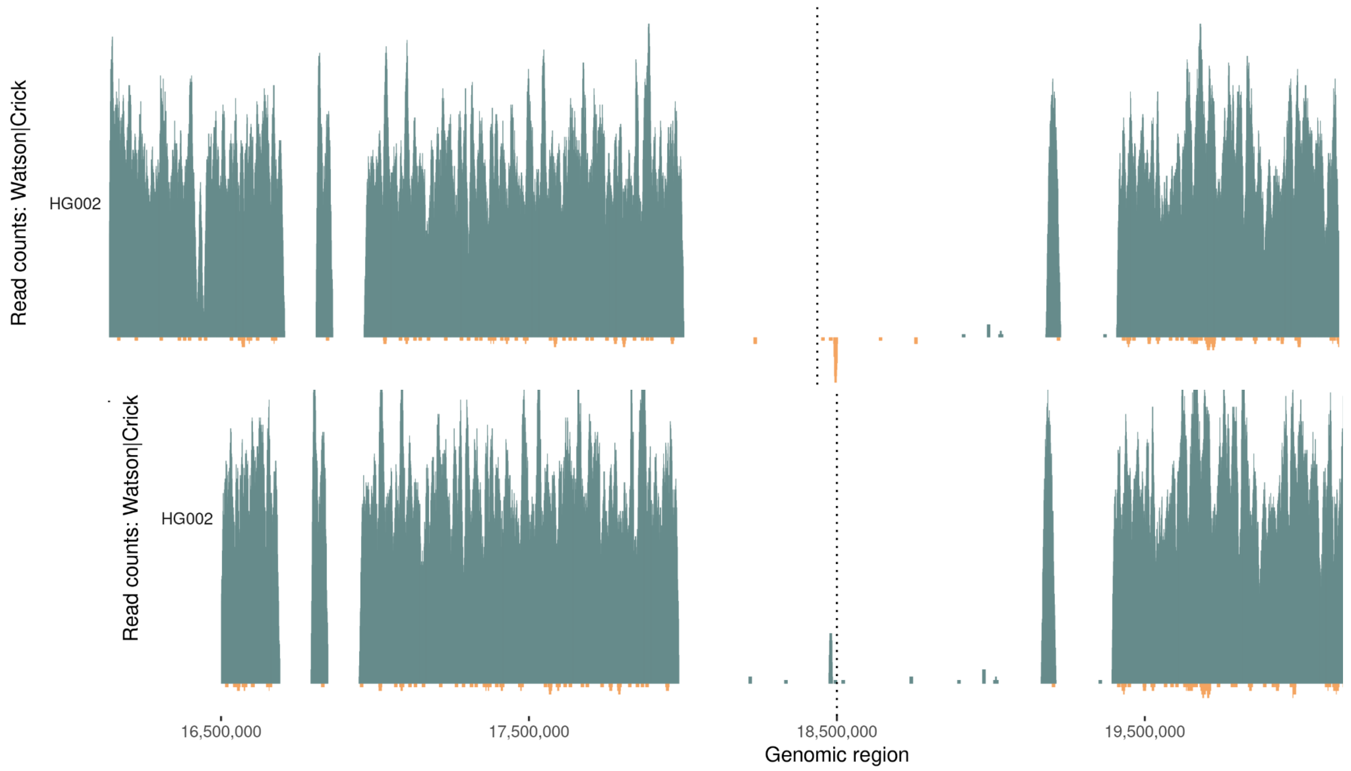


**Supplementary Figure 18.** Strand-seq alignments to the reference Chromosome Y before it was corrected (top) and full-coverage Verkko trio Chromosome Y assembly (bottom). Each plot shows Strand-seq directional read coverage reported as binned (bin size: 10,000, step size: 1,000) read counts represented as vertical bars above (teal; Crick read counts) and below (orange; Watson read counts) the midline.The top plot shows an inversion (dashed line) where directly oriented reads (Crick; teal) switch to inversely oriented reads (Watson, orange) and then back to directly oriented reads. The Verkko assembly in contrast is consistent with only Watson reads present in the same location (dashed line).

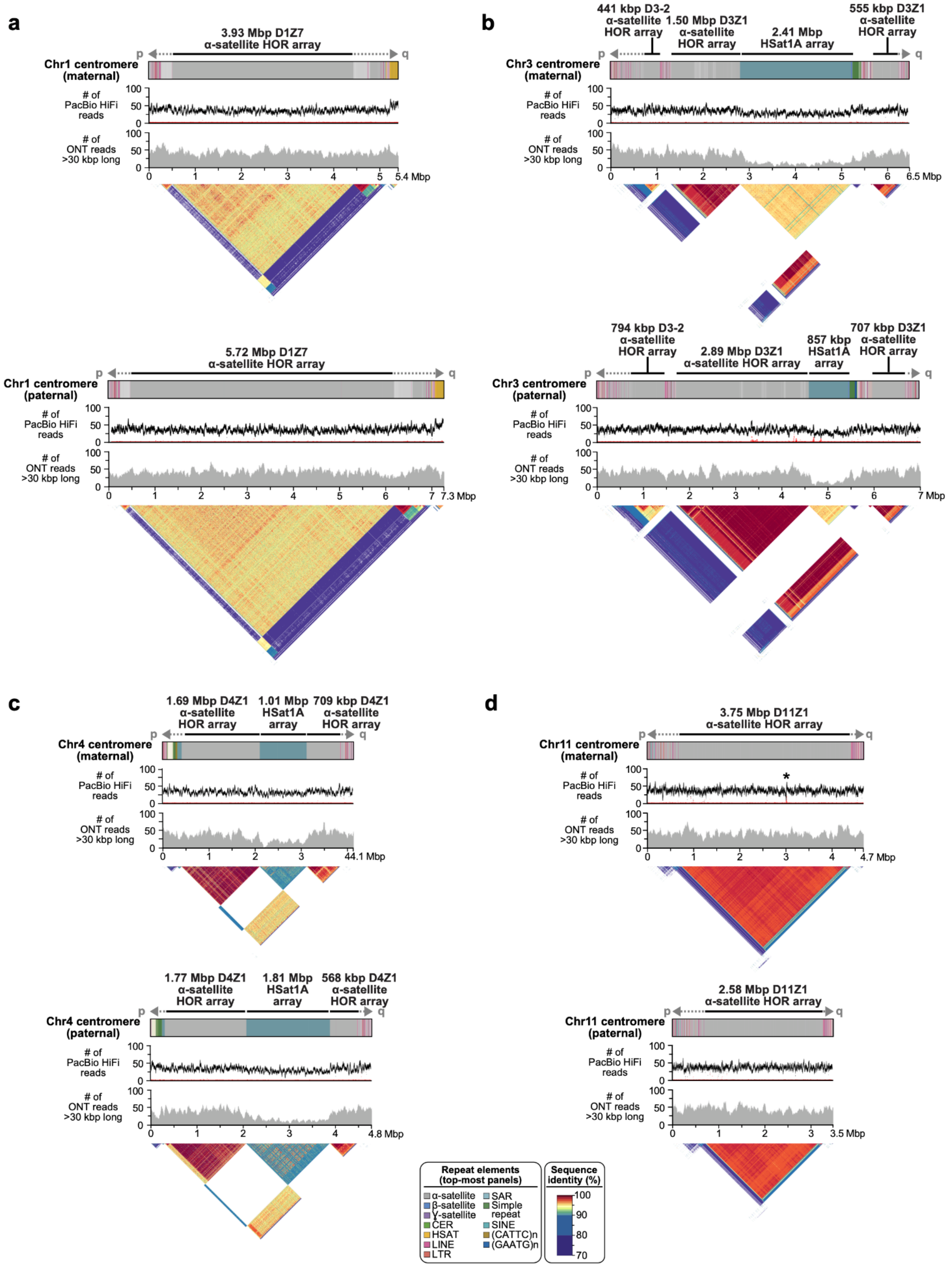


**Supplementary Figure 19.** Full-coverage Verkko trio assemblies of chromosome 1, 3, 4, and 11 centromeric regions in the HG002 genome. Both maternal and paternal haplotypes are shown, with repeat element annotation shown on top, followed by PacBio HiFi coverage, ONT coverage, and StainedGlass [58] plots. As with the Chromosome 19 centromeres (Fig. 4), the maternal and paternal haplotypes show large-scale structural variation, with alpha-satellite HOR arrays sizes varying by tens to hundreds of kb. Sites with discrepant HiFi mappings (low coverage or high coverage) are marked with an asterisk. There are few sites in the centromeres, and the artifacts are localized and often inconsistent between ONT and HiFi alignments, indicating the assembly is overall of high quality. To further validate assembly accuracy, we intersected centromere array locations with VerityMap errors and found that in all but two cases (both on the Chr1 paternal centromere), the errors were short (≤1 kb) or lower frequency (≤50% of the reads). VerityMap also identified one issue, with ≥50% of reads deviating in the Chr4 maternal centromere. However, this was not visible in the NucFreq [39,77] plots above, and the region only had a total of three mapped reads.

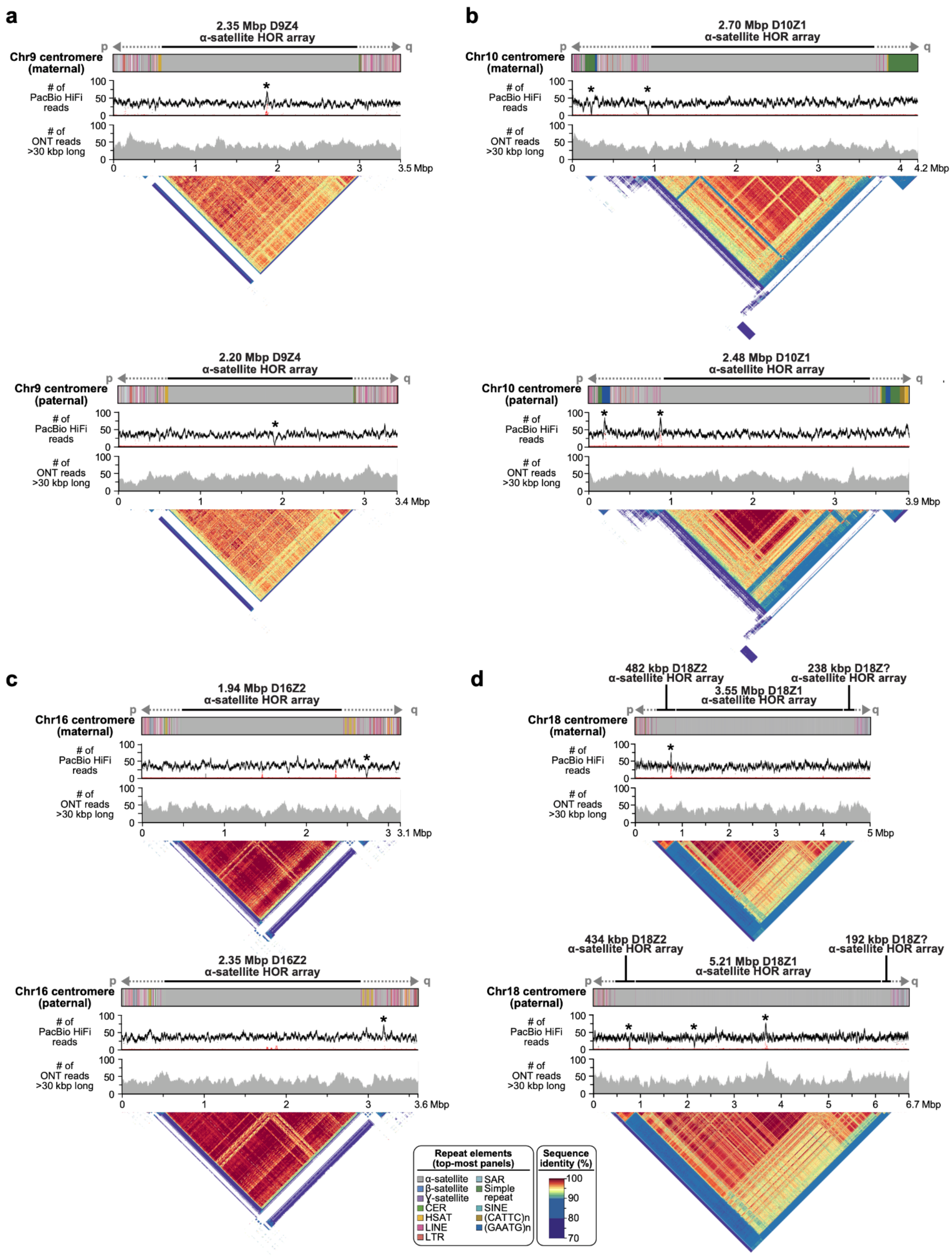


**Supplementary Figure 20.** Full-coverage Verkko trio assemblies of chromosome 9, 10, 16, and 18 centromeric regions in the HG002 genome. Both maternal and paternal haplotypes are shown, with repeat element annotation shown on top, followed by PacBio HiFi coverage, ONT coverage, and StainedGlass [58] plots. As with the Chromosome 19 centromeres (Fig. 4), the maternal and paternal haplotypes show large-scale structural variation, with alpha-satellite HOR arrays sizes varying by tens to hundreds of kb. Sites with discrepant

HiFi mappings (low coverage or high coverage) are marked with an asterisk. There are few sites in the centromeres, and the artifacts are localized and often inconsistent between ONT and HiFi alignments, indicating the assembly is overall high quality. To further validate assembly accuracy, we intersected centromere array locations with VerityMap errors and found that in all but two cases (Chr9 paternal centromere and Chr10 maternal centromere), the errors were short (≤1 kb) or lower frequency (≤50% of the reads).

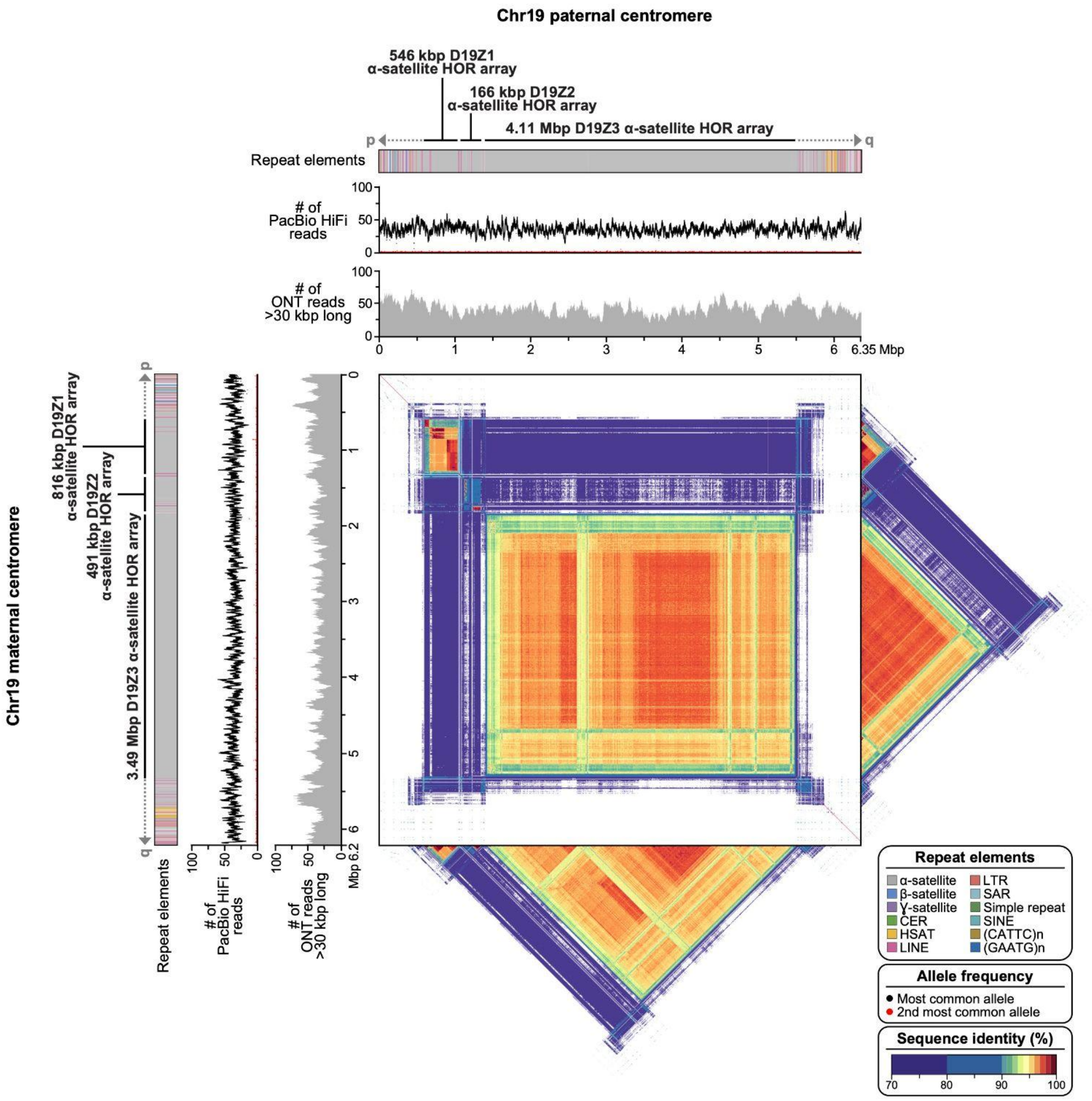


**Supplementary Figure 21.** Comparison of the HG002 maternal and paternal full-coverage Verkko trio assemblies for the centromeric regions of Chromosome 19. As in Figure 4, the tracks show the repeat annotations and read coverages. The central plot shows the similarity between the two haplotypes, with the maternal haplotype on the y-axis and the paternal on the x-axis. The triangles show the self-similarity within each haplotype for comparison.

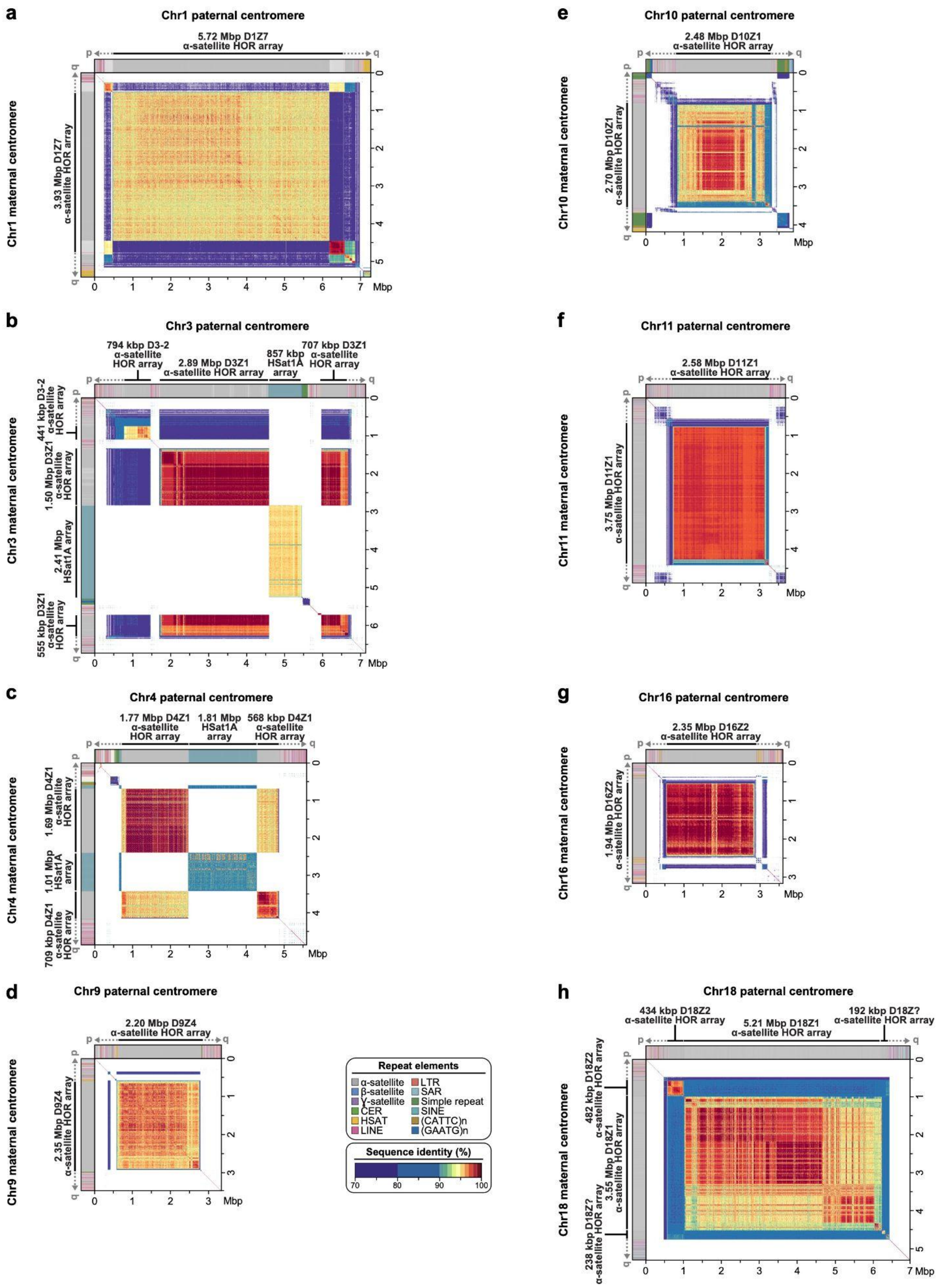


**Supplementary Figure 22.** Comparison of the HG002 maternal and paternal full-coverage Verkko trio assemblies for the centromeric regions of chromosomes 1, 3, 4, 9, 10, 11, 16, and 18 in the HG002 genome. The centromeric regions show varying α-satellite HOR array sizes and sequence identity between the two haplotypes, consistent with earlier reports that indicate that centromeric HOR arrays often expand and contract due to their repetitive nature and their propensity for unequal crossing over [78–80] and gene conversion [81] events.

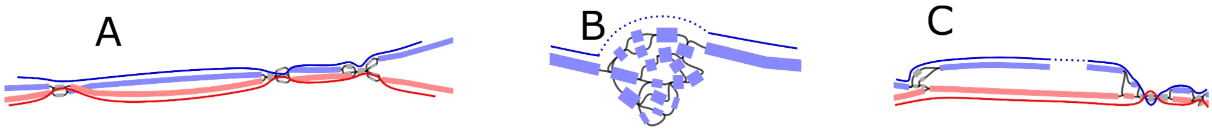


**Supplementary Figure 23.** Examples of haplotype scaffolding by Rukki in the HG002 genome. The nodes are colored according to their haplotype assignments. Nodes with at least 100 total markers where 90% of the markers agree are colored: red for maternal, blue for paternal. Nodes with less than 100 markers are colored gray for unassigned. The haplotype paths are marked with solid curves with dotted curves for gaps. (A) A well behaved genomic region consisting of phased heterozygous bubbles, homozygous nodes, and spurious nodes caused by sequencing errors. Where possible, Rukki connects the nodes attributed to the same haplotype across the homozygous regions, producing two phased unitigs without gaps. (B) A tangle within one haplotype. Rukki scaffolds across the tangle (dotted line), reporting an estimated size of the tangled region. (C) A gap in the paternal haplotype. Rukki uses haplotype assignments and the topology of the graph to scaffold across the gap (dotted line), and estimates the size of the gap based on the size of the paired haplotype.
